# Multiscale Modeling of Hepatitis B Virus Capsid Assembly and its Dimorphism

**DOI:** 10.1101/2022.02.23.481637

**Authors:** Farzaneh Mohajerani, Botond Tyukodi, Christopher J. Schlicksup, Jodi A. Hadden-Perilla, Adam Zlotnick, Michael F. Hagan

## Abstract

Hepatitis B Virus (HBV) is an endemic, chronic virus that leads to 800,000 deaths per year. Central to the HBV lifecycle, the viral core has a protein capsid assembled from many copies of a single protein. The capsid protein adopts different (quasi-equivalent) conformations to form icosahedral capsids containing 180 or 240 proteins, *T*=3 or *T*=4 respectively in Caspar-Klug nomenclature. HBV capsid assembly has become an important target for new antivirals; nonetheless the assembly pathways and mechanisms that control HBV dimorphism remain unclear. We describe computer simulations of HBV assembly, using a coarse-grained model that has parameters learned from all-atom molecular dynamics simulations of a complete HBV capsid, and yet is computationally tractable. Dynamical simulations with the resulting model reproduce experimental observations of HBV assembly pathways and products. By constructing Markov state models and employing transition path theory, we identify pathways leading to *T*=3, *T*=4, and other experimentally observed capsid morphologies. The analysis identifies factors that control this polymorphism, in particular, the conformational free energy landscape of the capsid proteins and their interactions.

## INTRODUCTION

During the lifecycles of many viruses, hundreds of protein subunits self-assemble into a protein capsid that packages the viral nucleic acid and delivers it to a new host cell. In many cases a particular capsid structure is required for infectivity, and capsid proteins form this structure with high fidelity. Yet, capsid proteins can also exhibit striking polymorphism and adaptability, assembling into structures with different sizes and morphologies in response to changes in solution conditions, the presence of antiviral drugs, or to encapsulate nucleic acids or nanoparticles of varying sizes. Elucidating the factors that control assembly pathways and products could identify important targets for antiviral drugs and would advance our fundamental understanding of the viral lifecycle.

Approximately half of characterized virus families have icosahedral symmetry. That is, their capsids are comprised of 60 identical asymmetric units. Caspar and Klug (CK) showed how multiples of 60 proteins can form icosahedral capsids, by sub-triangulating each asymmetric unit so that individual proteins are forced to adopt slightly different (quasi-equivalent) conformations [3–6]. An icosahedral capsid has 60*T* subunits, where the ‘triangulation number’ *T* specifies the number of different protein conformations, and is restricted to certain integer values (*T* = *h*^2^ + *hk* + *k*^2^, where *h, k* are non-negative integers) [4].

Although RNA sequences can drive allosteric switching of conformations in MS2 and closely related RNA bacteriophages [7–10], how the spatial arrangement of quasi-equivalent conformations is chosen during assembly is poorly understood for most viruses.

Hepatitis B virus (HBV) is an important example of self-assembly as a model system, and since chronic HBV infection is a serious public health issue, assembly of its capsid is a target for a new generation of antivirals that may contribute to a cure [11]. About 300 million people have chronic HBV, which contributes, by cirrhosis, liver failure, and liver cancer to about 800,000 deaths each year [12]. Within an infected host cell, the HBV capsid protein assembles around the HBV pre-genomic RNA and viral polymerase to form a core particle. While capsid structures with *T*=4 symmetry (comprising 120 dimer protein subunits) are appropriate to accommodate the genome, many secreted particles are empty (containing no RNA) [13] and a small fraction (~ 5%) have *T*=3 capsids (with 90 dimers) [14, 15]. This behavior can be recapitulated in vitro using the assembly domain (residues 1-149 of the capsid protein, Cp149) in the absence of nucleic acids, which leads to a similar mix of *T*=4 and *T*=3 particles (all of which are empty). Moreover, several classes of small molecule antiviral agents referred to as “Core protein Allosteric Modulators” (CpAMs) have been identified that can bind to HBV capsid proteins and drive inappropriate assembly leading to failure to package RNA or redirection of assembly to alternative noninfectious products [16–25].

Recently, Cp149 assembly size distributions have been measured near or at single-subunit precision using resistive pulse sensing [26, 27], mass spectrometry [28], and charge detection mass spectrometry [29–32], and small intermediates were identified by highspeed AFM [33]. These experiments identified a number of long-lived intermediate structures with sizes between 85-140 dimers. Notably some pathways also include ‘overgrown’ intermediates with more than 120 dimers, which eventually dissociate some dimers and rearrange into icosahedral capsids [34]. Similarly, small angle X-ray scattering (SAXS) experiments identified long-lived complexes at sizes between 90 and 120 dimers [35], as well as a three-phase assembly kinetics, with nucleation and elongation phases followed by a slow rearrangement of capsid subunits into icosahedral structures [36]. Despite the unprecedented detail enabled by these experimental advances, the mechanisms underlying HBV dimorphism and the factors that separate assembly pathways leading to *T*=4, *T*=3, or asymmetric products remain unclear.

Computational models can reveal details of the assembly process that are not accessible to experiments. However, while atomistic and near-atomistic simulations have revealed the dynamics of complete viruses [1, 37–43], the long timescales (ms-hours) prohibit simulating capsid assembly with atomic-resolution, except for specific steps [44, 45]. Therefore, coarsegrained models have been used to study the assembly dynamics of icosahedral shells[8, 46–90]. Of particular relevance to our work, elastic interactions between five-fold defects can funnel the elastic energy landscape of an assembling crystalline shell toward icosahedral structures[89–92]. However, previous works on empty capsids with (*T* ≥ 4) have focused on material properties (Young’s modulus and bending modulus) that are orders of magnitude different from those of virus capsids. Thus, different mechanisms may be important for HBV assembly.

A significant challenge of coarse-grained (CG) models is linking their predictions to specific experimental systems. It is therefore desirable to inform CG models with atomic-resolution information. For example, multiscale modeling approaches have elucidated the stability and dynamics of viral capsids [37, 40, 93, 94] and other protein structures (e.g. [42, 43, 95–111]), and CG models based on protein structures have led to insights about specific virus assembly processes (e.g. [112–118]).

Here, we employ data from atomic-resolution simulations and solution experiments to constrain and build a multiscale model for HBV capsid assembly. We start from a minimal CG model that captures the geometric features of HBV capsid proteins and their quasiequivalent conformations. We parameterize this model using data from all-atom molecular (AA) dynamic simulations of a complete HBV capsid, along with atomic-resolution structures of the proteins in different assembly morphologies. We focus on a small number of parameters: binding affinity, subunit length, subunit dihedral angle, binding angles, and associated stretching and bending moduli. This data provides estimates for the relative strengths of the different types of proteinprotein interactions that drive HBV assembly, the fluctuations of proteins around their mean interaction geometries, and the corresponding larger-scale elastic properties of the assembled structures.

We then perform dynamical Monte Carlo simulations to simulate dynamical assembly trajectories, and compare these to experimentally observed trajectories. Analyzing these computed trajectories using Markov state model and transition path theory analysis explains the origins of trapped or metastable species observed in experiments [29–32, 35, 36, 119]. These results provide the confidence to examine the early steps of the reaction, where the diversity of intermediates shows how nucleation can be described as a much more complex series of intermediates than could possibly be isolated in experiments. We identify pathway ‘hubs’, or intermediates from which pathways diverge toward *T*=4, *T*=3, or asymmetric assembly products. Pathway analysis isolates factors, such as the local symmetry and relative stability of associating subunits, that select which pathway emerges from a hub state. Further, for bending modulus values relevant to virus capsids (~ 40 - 400*k*_B_*T*), the elastic strain cannot guide assembly toward large symmetric structures, and robust assembly of icosahedral capsids with sizes *T* ≥ 4 strongly depends on the the presence of multiple protein conformations and their conformational dependant binding affinities. We find that such specificity of interactions plays a key role in the assembly of natural homopolymeric icosahedral capsids by reducing the number of accessible assembly pathways, and thus, guiding assembly toward the target geometry. Finally, the results in the paper are qualitatively applicable to other viruses, and the approach is generalizable.

## RESULTS

### Overview of model and parameters

Capsid protein dimers are the basic assembly subunits for HBV[120, 121]. Based on the quasiequivalent conformations of the monomers (denoted as A, B, C, and D), there are two dimer conformations in the *T*=4 capsid (AB and CD, see Fig. 1(A)), which each make four interdimer interfaces. Dimer-dimer binding is primarily driven by hydrophobic stereospecific contacts at these interfaces. In each such interaction, a monomer of one dimer is capped by a monomer of a second dimer, with the second fitting against a hydrophobic patch on the first. For example, in the D-B contact shown in Fig. 1(A), the D monomer in the CD dimer is capped by the B monomer in the AB dimer. These interactions are asymmetric — the inverse, a B-D contact with B of an AB dimer capped by D of a CD dimer, is structurally different from a D-B contact and, thus, has a different binding affinity.

**FIG. 1.**
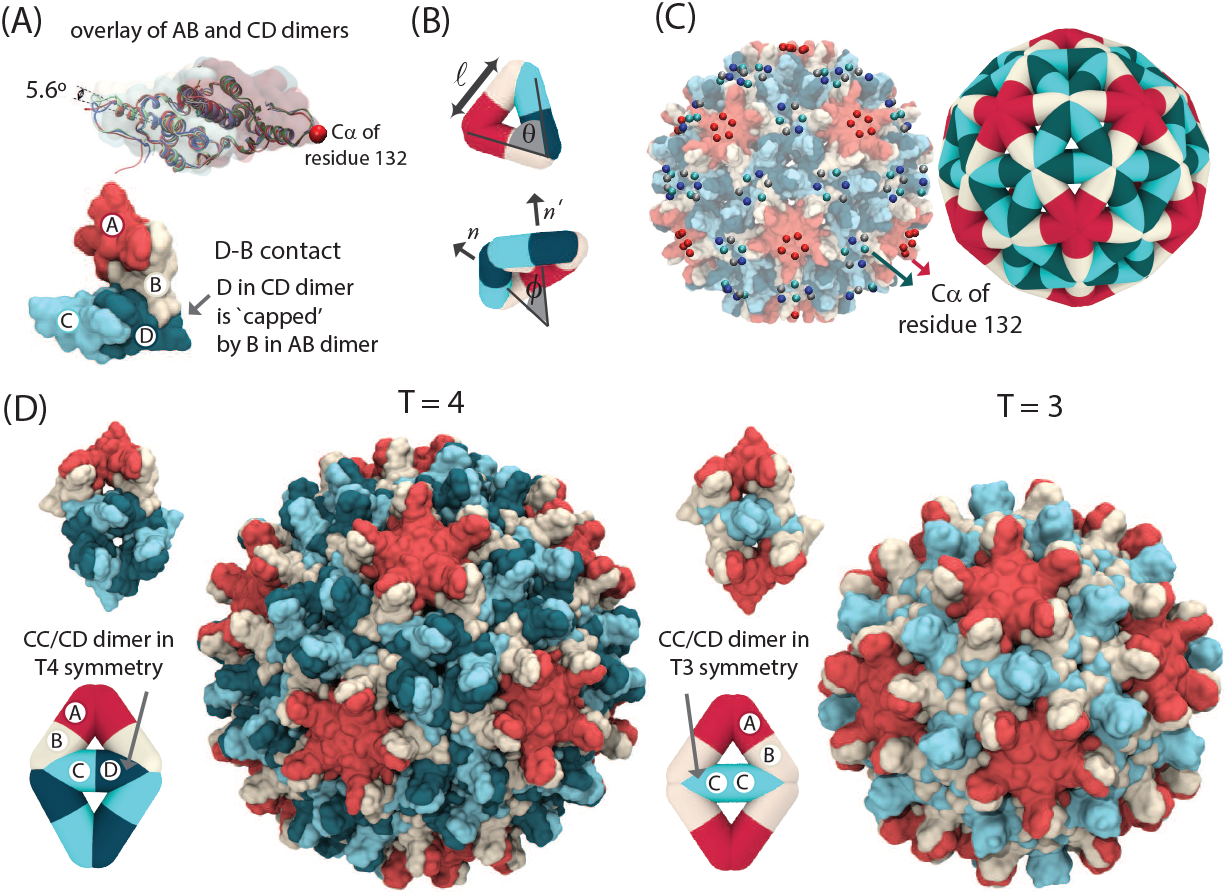
Description of the multiscale model. **(A)** Overlay of AB and CD conformations of the capsid protein from the crystal structure of the *T*=4 HBV capsid (top). Example of a dimer-dimer contact in the *T*=4 HBV capsid, where the D monomer in a CD dimer is ‘capped’ by a B monomer in an AB dimer (bottom). Colors correspond to different monomers in each dimer. **(B)** Each edge in the elastic sheet coarse-grained model represents an HBV dimer. Model parameters include the equilibrium edge length 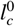 and associated force constant *κ*_l_, which controls the shell stretching modulus; the equilibrium binding angle between two edges 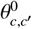 and associated force constant *κ_θ_*; and the equilibrium dihedral angle between the normals of the two adjacent triangles 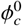 and force constant *κ_ϕ_*, which sets the capsid bending modulus. Here *c* and *c*′ are the conformations of a given edge and the dimer that it interacts with respectively. **(C)** Model parameters are optimized by minimizing the difference between the probability distributions of fluctuations of edge lengths, edge-edge angles, and dihedral angles computed from the coarse-grained (CG) model and an all-atom (AA) molecular dynamics simulation of a complete T=4 HBV capsid in explicit water [1]. We coarse-grain the data from the all-atom simulation by selecting the C-α atom of residue 132 of each monomer and clustering these 240 points to five-fold and six-fold vertices based on proximity using the scikit DBSCAN clustering algorithm [2]. The center-of-mass of each cluster is then assigned to a vertex of the CG model capsid (see SI Fig. S3 and movie S1). **(D)** The configurations of dimers in a *T*=4 capsid (left) and in a *T*=3 capsid (right). A CD/CC dimer in *T*=4 has two interactions with CD dimers and two interactions with AB dimers and is referred to a CD dimer. A CD/CC dimer in a *T*=3 capsid, which has 4 interactions with AB dimers, is referred to as CC dimer (see Model section and SI).

Since HBV dimers assemble with locally triangular lattices, we construct a CG model in which an assembling capsid is represented by an elastic triangular sheet, with edges corresponding to dimers. The approach is similar to existing models for capsid and tubule assembly based on triangular subunits of a single type [92, 122–124], but in our model growth occurs via addition/deletion of edges (dimers), and the model accounts for dimer asymmetry, as well as how the dimer structure and interactions depend on their conformational state (see the Model section, Fig. 1, and Fig. S1 of SI). We see below that these physical features are essential to achieve robust assembly of T=4 shells when accounting for the material properties of HBV proteins.

We consider two edge types with different equilibrium lengths and dihedral angles (Fig. 1(B) and Fig. S1). The first type represents AB dimers present in *T*=4 and *T*=3 capsids [125], while the second, CD/CC, represents both CD and CC dimers in *T*=4 and *T*=3 capsids respectively. This choice is based on the similarity of the structure and local symmetry environment (i.e. set of conformations of neighboring subunits) of CD and CC dimers in *T*=4 and *T*=3 capsids. For identifying assembly pathways, we will denote such a dimer as CD (CC) if its neighboring subunits are consistent with the *T*=4(*T*=3) local symmetry (see Fig. 1(D) and Fig. S1 of SI)

To model dilute, noninteracting capsid structures typical of productive capsid assembly reactions [58, 126], we perform dynamical grand canonical Monte Carlo (MC) simulations of a single assembling shell in exchange with free dimers at fixed chemical potential *μ*, which sets the bulk dimer concentration [122, 124]. The dynamics, including dimer association, obeys microscopic reversibility. Technical details are provided in the Model section and SI section Model Details.

#### Optimizing CG model parameters against AA simulations estimates protein-protein interaction geometries and elastic moduli

The model potential energy function includes terms that represent subunit-subunit binding interactions and the elastic energy associated with deviations from the ground state sheet geometry, corresponding to harmonic potentials for stretching of edges, deviations of dimer-dimer binding angles, and intra-dimer strains associated with deviation of dihedral angles between pairs of triangular faces (see Methods and Eq. (2)). Denoting a dimer conformation for edge *i* as *c*(*i*), the key model parameters are the set of equilibrium edge lengths 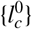, dimer-dimer binding angles, and dihedral angles 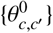, and 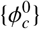 (Fig. 1(B)). The associated moduli, *κ*_l_, *κ_θ_*, and *κ_ϕ_* are independent of edge conformation, with *κ*_l_ and *κ_ϕ_* setting the continuum-limit stretching and bending moduli of the shell (see SI Model details).

To estimate the model parameter values that correspond to the HBV dimers, we mapped our coarsegrained dimers to an atomic-resolution structure of an HBV capsid (see methods and Fig. 1(C)). We then used AA simulations of a complete HBV capsid [1] to optimize the model parameters. We estimated the equilibrium edge lengths, dihedral and binding angles, and associated stretching and bending moduli of the CG model by minimizing the difference between the distributions of edge lengths, dihedral angles, and binding angles obtained from the AA MD and CG Monte Carlo simulations (see methods and Fig. S3). Optimization against the atomistic data resulted in high values for the stretching modulus *κ_l_* ≈ 4200*k*_B_*T*/*σ*^2^ (with the simulation unit length scale *σ* ≈ 8 nm) and binding angle modulus *κ_θ_* ≈ 800*k*_B_*T*, and a relatively low value for the bending modulus *κ_ϕ_* ≈ 40*k*_B_*T*. The 2D Young’s modulus *ϵ* ≈ 6000 can be estimated from 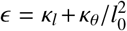, where *l*_0_ is the average dimer length ≈ *σ*.

Mapping the three bending and elastic moduli to the standard Helfrich elastic energy of an elastic sheet [127, 128], estimates the Föpple-von Kármán number, 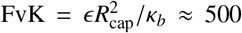 with *R*_cap_ ≈ 2.1*σ*, the radius of the HBV capsid, which is consistent with results of nanoindentation measurements [129]. As shown below, using these optimized elastic moduli in our model results in dynamical assembly behaviors that are consistent with experiments on HBV protein assembly; including the distribution of *T* =4 and *T*=3 icosahedral capsids at optimal assembly conditions, as well as large aberrant structures with reduced curvature observed at non-optimal conditions (including in the presence of CpAMs that strengthen dimer-dimer interactions [130]).

#### Binding affinities and conformational free energies

Estimates of the mean binding affinity (averaged over contacts between different dimer conformations and different interfaces) have been obtained from experiments that measure the yield of assembled capsids as a function of total subunit concentration for different solution conditions and temperatures [120]. These experiments identify a range of mean dimer-dimer binding affinity values for *T* =4 capsids from 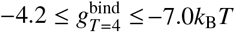 (with *k*_B_*T* ≈ 0.6 kcal/mol at *T* = 300K). To estimate the binding energies at each quasi-equivalent site, we set the CD binding affinity 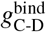 as a reference value 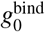, and set the relative values of binding affinities based on their relative buried surface areas (using PDBePISA, Table I). We define 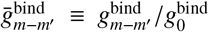 with *m, m′* as the conformations of the two interacting monomers. For example, 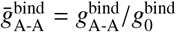, with analogous definitions for 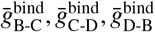, and 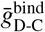. These estimated values result in simulated assembly behaviors that match those observed in experiments, whereas deviating from the PDBePISA estimates leads to poor agreement.

**TABLE I.**
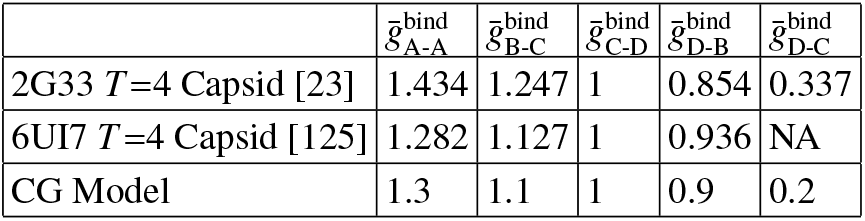
Relative dimer-dimer binding affinities. Relative binding affinities between different dimers in our CG model are set based on the inter-dimer interface analysis of HBV T = 4 capsid (in PDBePISA [153, 154]).

| | $\bar{g}_{\text{A-A}}^{\text{bind}}$ | $\bar{g}_{\text{B-C}}^{\text{bind}}$ | $\bar{g}_{\text{C-D}}^{\text{bind}}$ | $\bar{g}_{\text{D-B}}^{\text{bind}}$ | $\bar{g}_{\text{D-C}}^{\text{bind}}$ |
| --- | --- | --- | --- | --- | --- |
| 2G33 $T=4$ Capsid [23] | 1.434 | 1.247 | 1 | 0.854 | 0.337 |
| 6UI7 $T=4$ Capsid [125] | 1.282 | 1.127 | 1 | 0.936 | NA |
| CG Model | 1.3 | 1.1 | 1 | 0.9 | 0.2 |

However, we find that assembly morphology is very sensitive to the value of 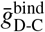. The D-C contacts do not occur within icosahedral structures, but are a prominent feature of flat hexagonal sheets that occur for nonassembly-competent conditions [131, 132] (Fig. 4(C-D)). Our simulations predict the optimal value for this parameter 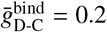 to be much less than the affinities of the contacts that are present in a *T*=4 capsid, and close to the PDBePISA estimate of ≈ 0.3

An additional unknown parameter is the equilibrium population distribution of the two dimer conformations AB and CD/CC. We specify this distribution according to 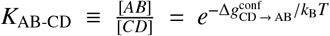, where *K*_AB-CD_ is the equilibrium constant for interconversion between the AB and CD conformations and 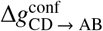 is the corresponding free energy difference between the two conformations. As noted above, we consider the CD and CC conformations to be equivalent, i.e. 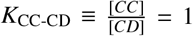, based on the high degree of structural similarity between CD and CC conformations and to reduce the number of model parameters.

To summarize, we consider two control parameters: 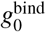 (which controls the mean inter-dimer binding affinity) and the intra-dimer conformtional free energy 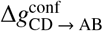. To enable direct comparison against experiments, we present our results in terms of the mean value of the dimer-dimer binding affinity (per dimer-dimer contact) in a *T*=4 capsid, which depends on the binding affinities and conformational free energy provided in Eq. 9 of the Model section. With the binding energy choices from Table I, the mean dimer-dimer binding affinity in Eq. 9 can be written as

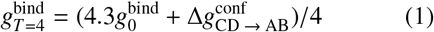

The mean binding energy 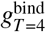 can be experimentally tuned by varying solution *p*H, ionic strength, or temperature as noted above [120]. The parameter 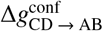 has not been directly estimated, but our results described below suggest that it depends on ionic strength, as previously suggested for interconversion of HBV capsid protein between conformations that are active or inactive for assembly [120, 133]. Further, experiments by Zhao et al. [134] suggest that 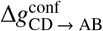 depends at least in part on entropic factors, such as different extents of disorder in the C-terminal region of the assembly domain (Cp149) for the different conformations. As described above, the other parameters are set according to atomic resolution simulations (bending and stretching moduli, equilibrium angles and edge lengths) or structures (relative strengths of binding affinities for different conformations) and are expected to be relatively insensitive to solution conditions. However, binding affinities for different conformations could be changed by amino acid substitutions [135] or small molecule binding at dimer-dimer interfaces, and thus we consider the effect of varying 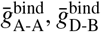, and 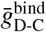 below.

### Assembly products

#### The model reproduces HBV polymorphism

Fig. 2(A-C) shows how the distribution of assembly products depends on the mean dimer-dimer binding energy 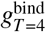 and the AB/CD conformational free energy difference 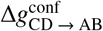. We classify the assembly products into three categories (with representative snapshots shown in Fig. 2(D)): *T*=4 capsids; *T*=3 capsids; and *malformed* structures, which do not have icosahedral symmetry but are (meta)stable on simulation timescales. Fig. 2(A) shows the selectivity for *T* =4 capsids (defined as ratio of *T*=4 capsids to all closed shells) as a function of 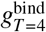 and 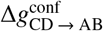, while Fig. 2(B-C) shows histograms of assembly product distributions for varying 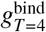 and 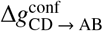 respectively.

**FIG. 2.**
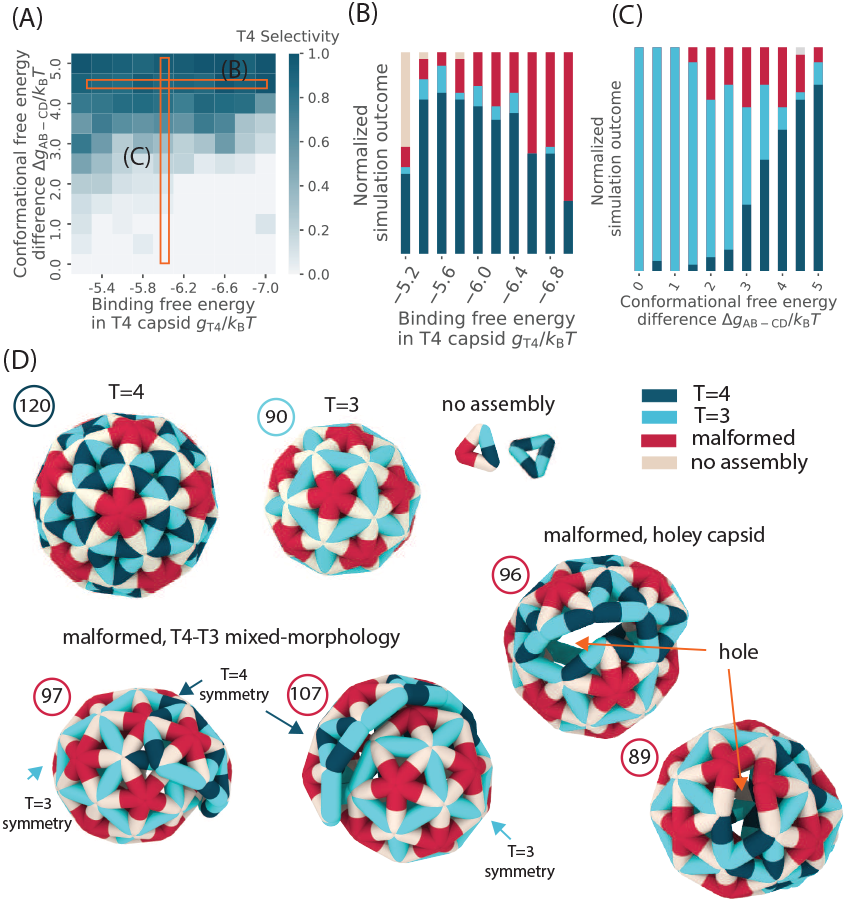
Dependence of assembly product morphologies on binding affinities and the intra-dimer conformational free energy landscape. **(A)** Dependence of selectivity for *T*=4 capsids on the mean dimer-dimer binding affinity in a *T*=4 capsid (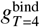, Eq. (1)) and the conformational equilibrium between AB and CD dimers, parameterized by 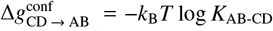. The selectivity is defined as the ratio of the number of complete *T*=4 capsids to the number of all closed shells. **(B,C)** The fraction of assembly product morphologies as a function of **(B)** the dimer-dimer binding affinity for fixed 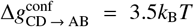 and **(C)** the AB/CD conformational equilibrium for fixed 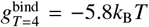. The four categories of product morphologies are shown at the top. Capsids with *T*=4 and *T*=3 icosahedral symmetry have 120 and 90 dimers, respectively, while most shells with mixed morphology have sizes between 85 and 140 dimers. **(D)** Snapshots of the three categories of assembly products are shown; *T*=4, *T*=3 and long-lived mixed-morphology capsids that have two distinct parts in *T*=4 and *T*=3 symmetries. Long-lived mixed morphologies (left) fail to close due to incompatible curvatures of the two morphologies, resulting in open boundaries that are sterically blocked from addition of new dimers (see SI Fig. S4). Holey capsids (right) frequently form under aggressive assembly conditions, when the two capsid regions with different symmetries bind imperfectly.

The model reproduces the experimental observations that a fraction of *T*=3 capsids assemble, despite the fact that the model is geometrically optimized for the *T* =4 icosahedral symmetry. Further, the malformed products have a broad size distribution with most shells between ~ 85 – 140 dimers, which is consistent with size distributions measure by CDMS [31]. Malformed structures are typically unclosed, with a mixed morphology that comprises two distinct parts with *T* =4 and *T*=3 morphologies, although we observe strained holey capsids at high binding energies. Examples of mixed-morphology and holey capsids are shown in Fig. 2(D).

#### The intra-dimer conformational equilibrium strongly affects the ratio of *T*=4/*T*=3 capsids

As shown in Fig. 2A, the model robustly assembles icosahedral capsids over a broad range of 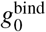 and 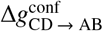 but the selectivity for *T*=4 capsids depends strongly on the conformational free energy. For a relatively narrow range 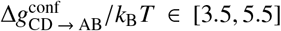, we observe selectivity values that are consistent with experimental observations (on the order of 5-30%). This suggests bounds for the parameter 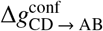.

#### Overly strong binding affinities increase the fraction of malformed structures

Increasing the base binding affinity 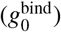 decreases the proportion of assembled shells that have *T*=4 symmetry (x-axis of Fig. 2(A)). Interestingly, the yield of *T*=3 capsids is relatively constant over this range as observed in [136]; the reduction in *T* =4 morphologies occurs due to an increase in malformed structures, especially the mixed-morphology class. These structures typically have open boundaries, and in moderate assembly conditions (moderate binding affinities or subunit concentrations) they are able to ‘edit’, or reconfigure their assembly geometry to result in icosahedral *T*=3 or *T* =4 capsids (e.g. Fig. 5(B) below). We discuss the pathways and mechanisms leading to such mixed-morphology structures below.

#### Simulated assembly product size distributions are consistent with experiments if the intradimer conformational equilibrium depends on ionic strength

Fig. 3(A) shows the assembly product size distribution measured in CDMS experiments at four different parameter sets, corresponding to two subunit concentrations and two ionic strengths, from Ref. [31]. The fraction of *T*=3 capsids dramatically decreases with ionic strength, from ~ 5% at *I* = 210 mM (top) to ~ 30% at *I* = 510 mM (bottom). In contrast, increasing the subunit concentration from *C*_tot_ = 10 *μ*M (left) to 20 *μ*M (right) does not significantly change the *T*=4/*T*=3 ratio, but does increase the prevalence of non-icosahedral structures with sizes between 85-140 dimers.

**FIG. 3.**
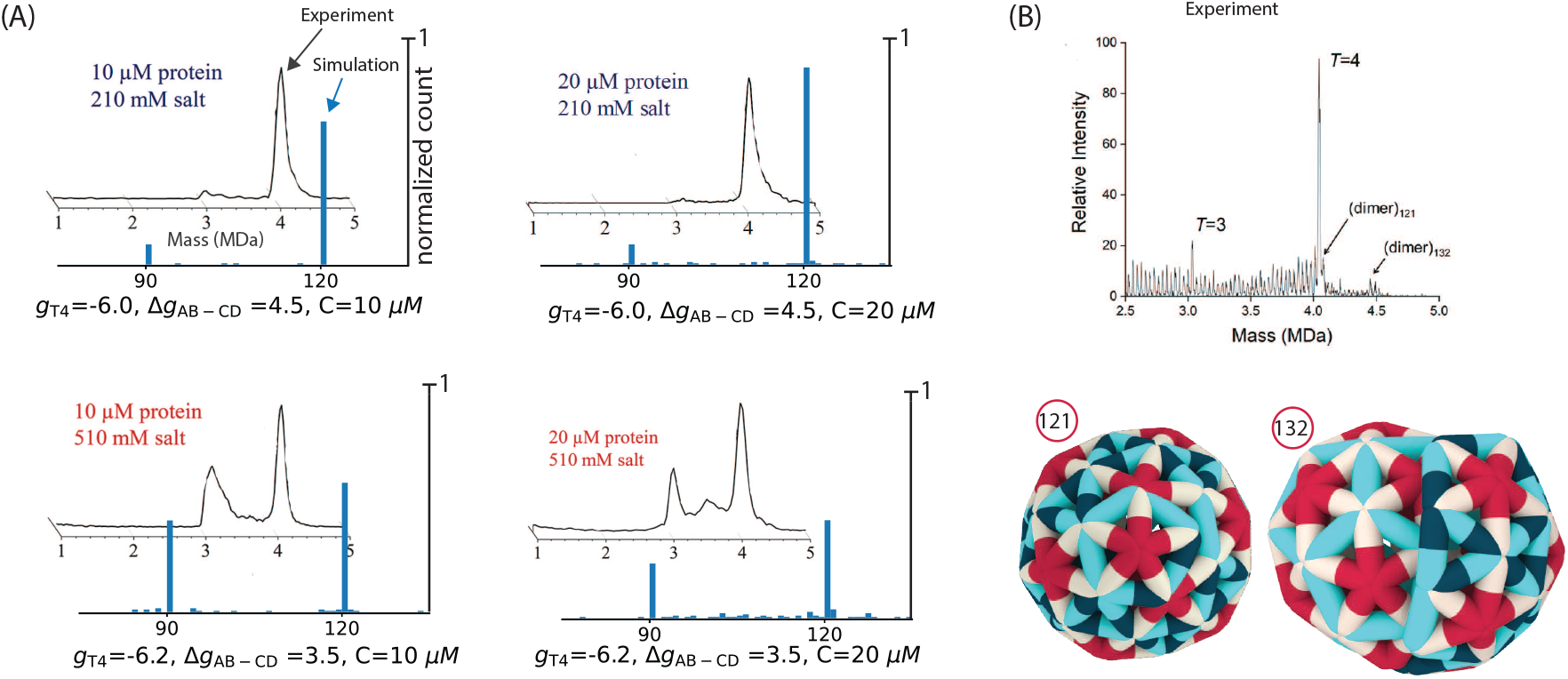
**(A)** Comparison of the simulation assembly product size-distribution with CDMS experiments (after 10 minutes). Experimental results are shown for ionic strengths of *I* = 210 mM (top) and *I* = 510 mM (bottom); and dimer concentrations *C* = 10 *μ***M** (left) and *C* = 20 *μ***M** (right). The peaks near 3 and 4 MDa correspond to *T*=3 and *T*=4 capsids respectively. Simulation assembly size distributions are shown for 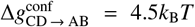, 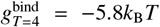 (top) and 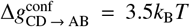, 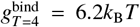 (bottom); and *C* = 10 *μ*M (left) and *C* = 20 *μ*M (right). Insets of CDMS spectra in the four panels are respectively from Figure 1 A-D of Lutomski et al. [34]. **(B)** High-resolution size distribution of assembly products in CDMS experiments [32] for *I* = 200 mM and *C* = 10 *μ*M. Snapshots show the mixed-morphology capsids with 121 and 132 dimers, which are among the most abundant non-icosahedral structures observed in both simulations and experiments. While the 121-dimer shell has 111 of its dimers consistent with *T*=4 symmetry and a small region of 10 dimers with local *T*=3 symmetry, the 132-dimer shell contains an approximately equal mix (≈ 60 dimers in *T*=3 and ≈ 62 dimers in *T*=4 environments).

We find that the simulations qualitatively reproduce both of these behaviors (simulation results in Fig. 3(A)) if the conformational free energy 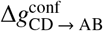 decreases (*K*_AB-CD_ increases) with increasing ionic strength. In particular, decreasing the free energy difference from 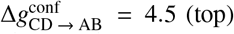 (top) to 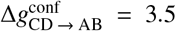 (bottom) changes the *T*=4/*T*=3 ratio, similar to the effect of increasing the salt concentration in experiments. Here we have also slightly increased the binding affinity, 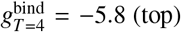 (top) and 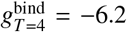 (bottom) to match the experimental observation that 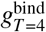 increases with increasing the salt concentration. Increasing the protein concentration (from left to right) results in more mixed-morphology products in simulations and CDMS experimental observations. These results are consistent with a previous suggestion of a relationship between dimer conformation and ionic strength based on a range of experimental observations [120, 133].

Fig. 3(B) shows a high-resolution CDMS measurement of the assembly product size distribution for *I* = 200mM and C = 10*μ*M. The assembly product size distribution in our higher concentration simulations are qualitatively consistent with the experimental observation, and we observe prevalent non-icosahedral assembly products at consistent sizes; e.g., the 121-dimer and 132-dimer structures shown in the figure.

#### The conformational dependence and asymmetry of HBV interdimer interactions are highly optimized

The results discussed to this point have focused on the relative values of dimer-dimer binding affinities estimated from buried surface area (PDBePISA) for each pair of dimer conformations (Table I). Since we expect these to roughly correspond to the wild type HBV capsid protein, changes in these affinities would correspond to amino acid substitutions at the dimer-dimer assembly interface, or the presence of antiviral agents that bind to the assembly interface [16–25]. Note that the dimer-dimer interactions are asymmetric — within a capsid each dimer is ‘capped’ from one side by another dimer, and thus, for example, the C-D contact (with binding energy 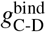) where C is capped by D is different from the D-C contact (with binding energy 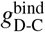) where D is capped by C.

Fig. 4(A-B) shows the assembly products that result when some of the relative binding affinities (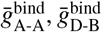, and 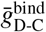) deviate from the PDBePISA estimate. Remarkably, these results suggest that the binding affinities are highly optimized for *T* =4 capsid assembly. Selectivities for *T*=4 on the order of those observed experimentally occur only for relative binding affinities that are close to those estimated from buried surface area (denoted by the ‘*’ symbols in Fig. 4(A-B)).

**FIG. 4.**
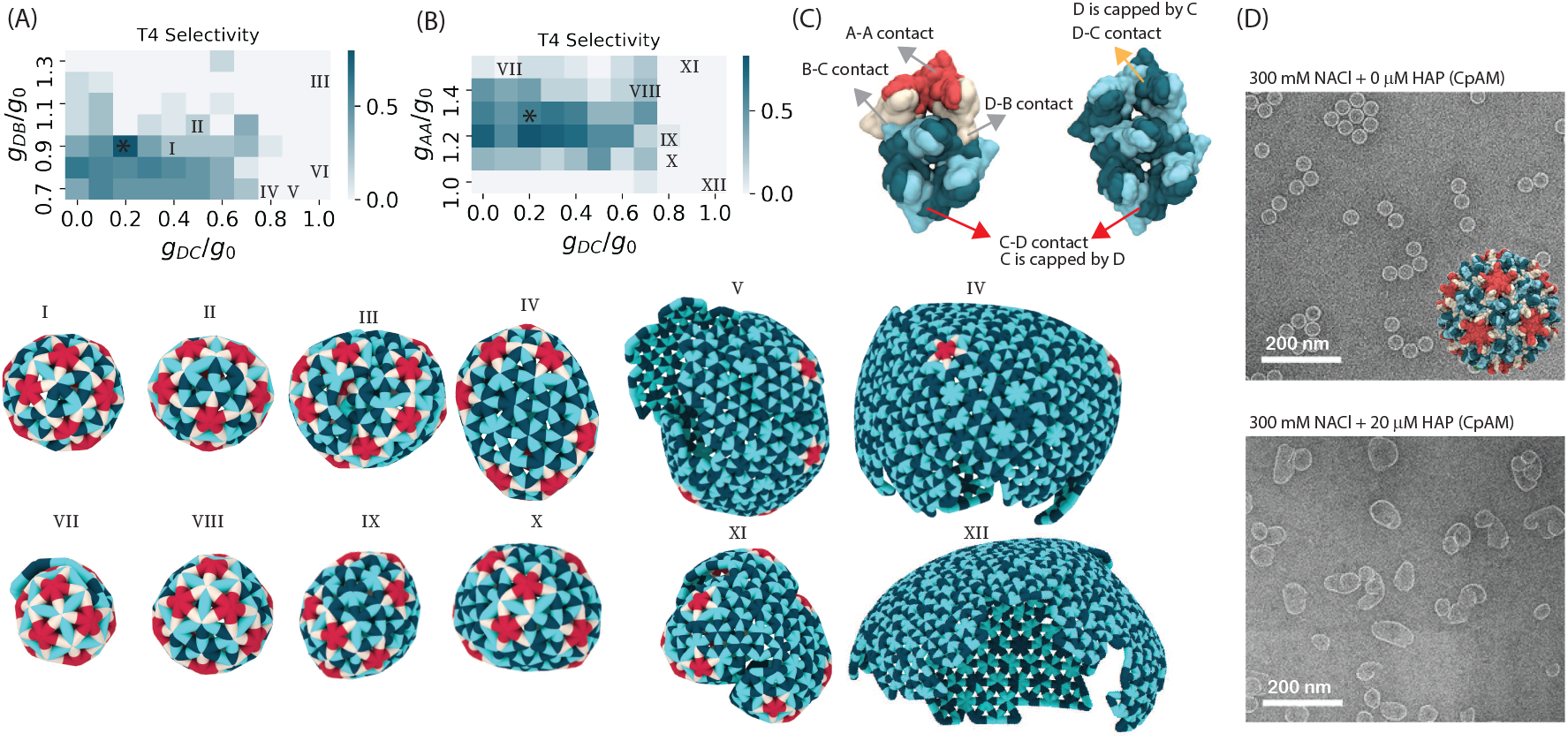
Conformational specificity of interdimer binding affinities is important for assembling *T*=4 capsid morphologies. **(A,B)** Dependence of selectivity for *T*=4 capsids on binding affinities between dimers with different conformations: **(A)** 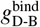 and 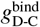; **(B)** 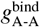 and 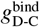. Selectivity is defined as the ratio of *T*=4 capsids to all closed shells. The ‘*’ symbols in (A) and (B) indicate the relative binding affinities estimated from PDBePISA, showing that these values result in *T*=4 selectivities close to those observed in experiments. Snapshots of typical morphologies at parameter sets indicated on the plots (I-XII) are shown at the bottom. **(C)** A diamond from a *T*=4 capsid (left) and a small piece of the hexameric lattice (right) that forms in flat sheets, showing different contacts between quasi-equivalent conformations. The D-C contact is required to form the hexameric sheets and aberrant structures, but does not occur in *T*=3 or *T*=4 morphologies. **(D)** Transition electron micrographs of the in vitro assembly of HBV dimers in the absence (top) and presence (bottom) of CpAM antiviral molecules (images adapted from [130]).

Deviations in 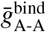 and 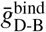 have a qualitatively different effects on assembly morphology than 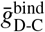. First, optimal values of 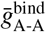 and 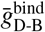 are ~ 1, whereas the optimal value of 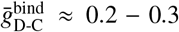 (i.e. the D-C contact is much weaker than the mean binding affinity). Second, increasing 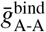 and 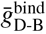 above their optimal values typically results in mixed-morphology malformed structures described above for overly strong mean binding affinities. In contrast, increasing 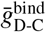 above its optimal value, 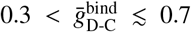, leads to assemblies that are closed but lack even partial icosahedral symmetry. In most cases these are asymmetric, but at some parameter sets we observe capsids with D5H symmetry (e.g. image I at 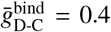 in Fig. 4). Further increasing 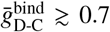 results in large aberrant structures that contain extended hexagonal lattices (V, VI, XI and XII in Fig. 4). Interestingly, these structures bear a strong resemblance to those observed in experiments on assembly of HBV dimers in the presence of core protein allosteric modulator (CpAM) molecules Fig. 4(E) [130]. CpAMs in general are assembly agonists, increasing dimer-dimer binding affinities; HAPs in particular favor a flattening of quasi-sixfolds by binding preferentially to B-C and C-D interdimer interfaces [20]. Increasing the D-C affinity qualitatively mimics this effect.

The strong dependence of assembly morphologies on conformational specificity can be attributed to the relatively low bending modulus estimated from the atomistic simulations (and consistent with experiments), *κ_ϕ_* ≈ 40*k*_B_*T*. The difference in elastic energy per-subunit between *T*=3 and *T*=4 morphologies is 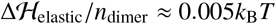, and even the large aberrant structures (V, VI, XI and XII in Figure 4) incur only modest elastic energy costs relative to a *T*=4 structure 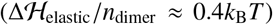. Thus, gradients in elastic energy alone are not strong enough to guide assembly trajectories toward the *T*=4 morphology, and conformational dependence is required as well.

For the remainder of this article, all results use the default set of values for 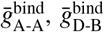, and 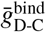 listed in Table I.

### Assembly pathways

#### Error correction during assembly and recovery from overgrown structures

Fig. 5 (A-D) shows example assembly trajectories that respectively result in *T*=3 or *T*=4 capsids, or asymmetric assembly products. Snapshots are shown along each trajectory, along with labels indicating the numbers of dimers in long-lived intermediates. Here we have defined a long-lived intermediate as a state with a population fraction (of the ensemble of all states along dynamical trajectories) that exceeds a threshold value of 1 % (Fig. S5(A) in SI). Qualitatively, this corresponds to the fraction of protein in detectable intermediates in SAXS [35, 137].

**FIG. 5.**
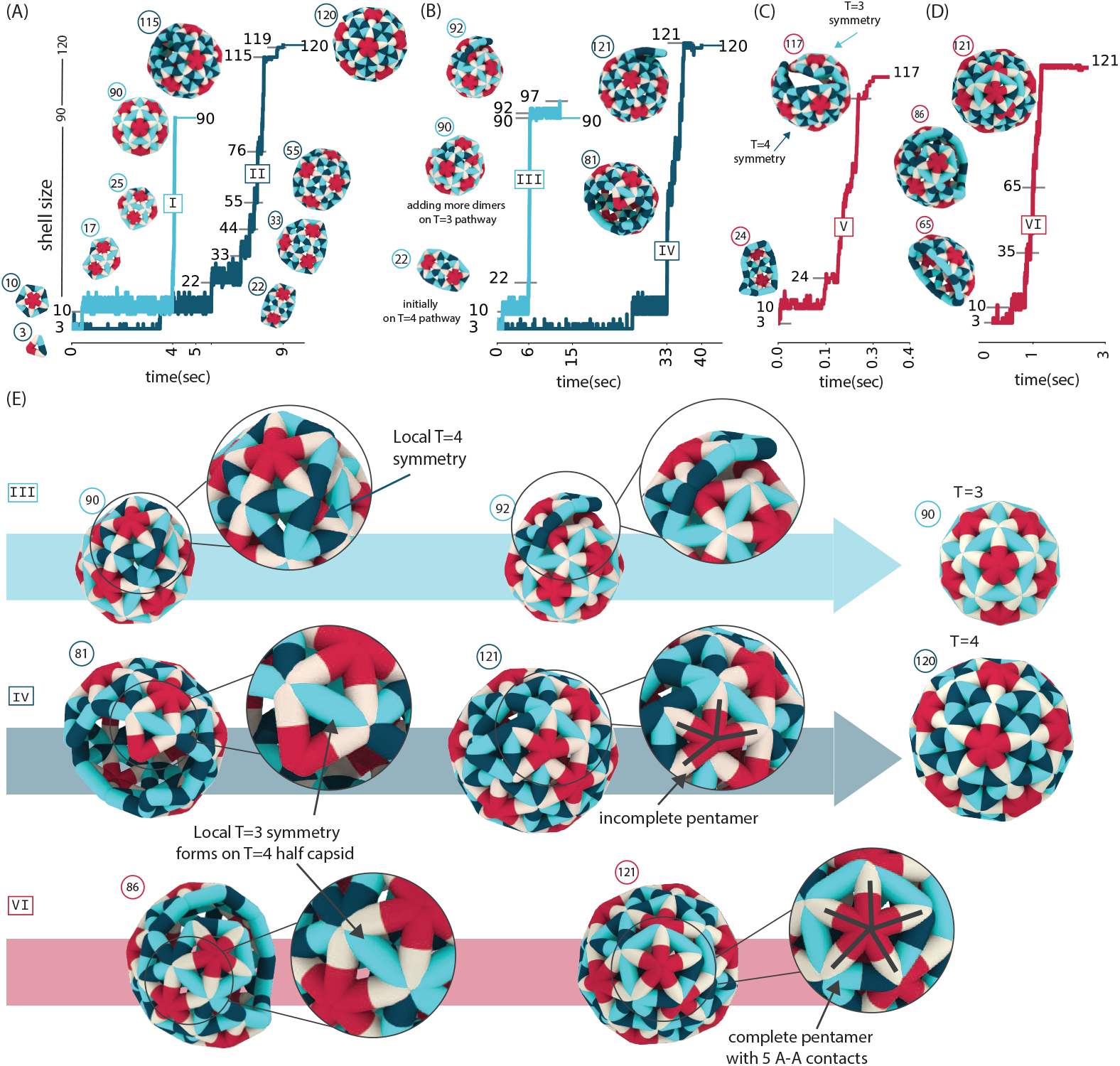
Assembly pathways. **(A)** Examples of assembly trajectories of *T*=4 and *T*=3 capsids, with snapshots of assembly intermediates at 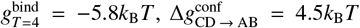, and *C* = 20*μ*M. **(B)** Examples of assembly trajectories in which overgrown (unclosed) capsids form, followed by ‘error correction’, or shedding of excess subunits and reconfiguration into complete *T*=4 (blue) and *T*=3 (cyan) capsids. Parameters for the *T*=4 and *T*=3 trajectories are respectively 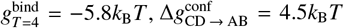, *C* = 15*μ*M; and 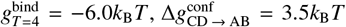, *C* = 10*μ*M. **(C)** Example trajectory leading to a mixed-morphology capsid with a size (117 dimers) between *T*=3 and *T*=4. Parameters are 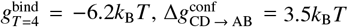, and *C* = 20*μ*M. **(D)** Example trajectory resulting in a long-lived mixed-morphology shell with 121-dimers. Parameters are 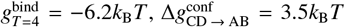, and *C* = 20*μ*M. (E) Zoomed-in views of assembly intermediates of trajectories in (A)-(D) show formation of locally incompatible morphologies, which are corrected in (A) and (B), but remain in the structure in (C) and (D) trajectories. If the dimers in a locally incompatible morphology end up in a closed pentamer [VI], correction is usually not observed on simulation timescales. Animations corresponding to these trajectories are provided in SI movies S2-S6.

While the assembly products can be classified into different categories as described above, the underlying assembly pathways have a key feature in common. At multiple points along assembly trajectories, subunits can bind with a local geometry that is incompatible with the larger-scale geometry of the existing shell, resulting in regions corresponding to different morphologies (e.g. a combination of *T*=4 and *T*=3 symmetries). This morphology mismatch results in different behaviors depending on the extent of the mismatch and the number of subunits required to dissociate or rearrange to achieve a uniform morphology.

*T*=4 assembly trajectories (Fig. 5(A)) typically involve several long-lived intermediates, which have geometries such that the next dimer to associate can only form a single bond and is thus relatively unstable. In addition, many of these states have several CD/CC dimers on the boundary, which can act as a ‘seed’ for mixed-morphology excursions in which dimers bind with local *T*=3 symmetry. In this case, successful assembly of a *T*=4 structure requires error correction, i.e., unbinding or reconfiguration of the dimers in local *T*=3 arrangements to recover global *T* =4 symmetry. Thus, trajectories may surmount multiple free energy barriers before assembly proceeds rapidly.

While both *T*=3 and *T*=4 assembly exhibit a very long-lived 10-dimer intermediate, we do not observe larger long-lived intermediates in *T*=3 pathways. This trend reflects the fact that the on-pathway *T*=3 intermediates with sizes *n* > 10 have either a convex boundary or multiple AB dimers, and thus avoid high barriers to additional subunit association or seeds with local *T*=4 symmetry. This suggests that the relatively high value of 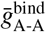 plays a key role in the facile completion of *T*=3 capsids.

##### Overgrown capsids

In some cases, regions with mismatched local symmetry remain in the partial capsid structure until the final stages of the assembly, resulting in the formation of ‘overgrown’ structures with more than 120 or 90 dimers for *T*=4 or *T*=3 capsids respectively. Fig. 5(B) shows an example of a trajectory (blue line) for which a region of local *T*=3 symmetry causes a mismatch between the orientations of boundary dimers as the capsid nears completion, resulting in overgrowth to an unclosed 121-dimer structure. However, eventually the dimers in the region with *T*=3 symmetry disassemble due to their less favorable elastic energy and binding affinities, and then the capsid rapidly forms a complete *T*=4 structure. We observe similar pathways involving correction of overgrown structures that lead to *T*=3 capsids. For example, Fig. 5(B) (cyan line) shows a trajectory in which an overgrown mixed-morphology structure with about 97 dimers eventually closes with *T*=3 symmetry. This pathway starts with formation of a 22-dimer structure in *T*=4 symmetry, and continues with addition of dimers in *T*=3 symmetry, until reaching a metastable intermediate with 90-92 dimers, that retains the initial *T* =4 region. This long-lived intermediate has ≈ 9 dimers on the boundary, but further growth is prevented by sterics and elastic strain due to the unfavorable curvature (see Fig. S4(A) of SI). Finally, the structure breaks interactions and undergoes conformational changes in the *T*=4 region, leading to the *T*=3 morphology.

The above-described trajectories and error correction are qualitatively consistent with the recent CDMS observations of overgrowth followed by shedding of excess subunits [34]. Moreover, slow capsid closure during the final stages of HBV assembly was observed in SAXS experiments by Chevreuil et al. [36].

In particular, they found that assembly preceded by three stages, nucleation, growth, and closure, with the closure phase involving a longer timescale than the growth phase. Figure S5(B) in SI shows that *T* =4 trajectories on average might spend up to ≈ 20% of the assembly time in the pre-closure states.

Importantly, such error correction only occurs under moderate assembly conditions; i.e., when the net driving force for assembly (determined by binding affinities and dimer concentrations) is such that assembly is nearly reversible. For example, the fraction of mixed-morphology assembly products increases with 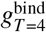 in Fig. 2(B), and we observe a similar dependence on total dimer concentration.

#### Long-lived off-pathway intermediates in simulations and experiments

At higher concentrations or higher binding affinities, we observe a variety of assembly products (meaning they are (meta)stable on all simulation timescales) with sizes between between 85-140 dimers. Fig. 5(C) shows an example trajectory that maintains *T* =4 symmetry until reaching 65 dimers, but then a local *T*=3 symmetry region forms and dominates the subsequent growth. The mismatch in curvature between the *T*=4 and *T*=3 regions leads to an unclosed structure. Despite an open boundary that allows further subunit association and dissociation, the size of the structure remains relatively constant at ≈ 117 dimers because the strain resulting from the curvature mismatch competes with the net assembly driving force. Because this structure does fluctuate in size, we anticipate that, if simulated long enough, it could eventually form a complete *T*=3 or *T* =4 shell. This speculated outcome would be consistent with experiments that observed intermediates with sizes between 85-140 dimers after several hours, but showed that on longer timescales (72 hours) most intermediates were converted into complete *T*=3 and *T* =4 capsids [32].

Fig. 5(D) shows another example trajectory, which results in the long-lived 121-dimer state. When the capsid is approximately half-complete, a region with local *T*=3 symmetry forms and then remains through-out the assembly process, Consequently the shell curvature is slightly distorted, which prevents the assembly from closing on itself. Instead, the assembly starts to spiral around itself before stalling at a 121 dimers, resulting in a ‘holey capsid’. Despite being incomplete, this structure is highly metastable because the dimers in the incorrect local symmetry are stabilized by the relatively strong BA-AB binding affinity 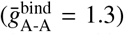, while addition of new dimers is sterically obstructed by the spikes of neighboring dimers (see Fig. S4(D) of SI).

Fig. 5(E) shows zoomed-in views of some overgrown structures. Comparing the two 121-dimer structures in Fig. 5(B) and Fig. 5(D) reveals an insight into the relative metastability of such states. While overgrowth results in both of these trajectories because a local *T*=3 forms within a predominantly *T* =4 shell, the *T*=3 regions in [IV] or [VI] respectively contain incomplete or complete pentamers. Consequently, the overgrown structure in [IV] sheds excess subunits and rearranges into the stable *T* =4 morphology relatively quickly, whereas that in [VI] remains metastable on simulated timescales. This behavior demonstrates that similar assembly ‘errors’ can have different effects on pathway selection and assembly products because small differences in local geometry become amplified as a structure grows. Thus, the sizes of different morphology regions within a structure affect its degree of metastability — a few incorrectly bound dimers with weak interactions can rearrange on experimentally relevant timescales, whereas larger regions of incompatibility cannot.

### Inferring mechanisms of dimorphism and path selection

Following the method introduced in Ref. [138] we built Markov State Models (MSMs) from the simulated assembly trajectories, and then used transition path theory [139] to enable further insight into factors that control morphology selection during assembly, and when pathways resulting in *T* =4 or *T*=3 morphologies diverge from each other. Constructing an MSM requires defining a state space and estimating the transition probability matrix between all pairs of states. To this end, we decomposed the set of simulated capsid structures into a state space defined by two order parameters: (1) the number of dimers *n*_dimer_ in a partial capsid structure, and (2) an order parameter that distinguishes the assembly morphology. We chose different quantities for the second order parameter depending on the morphology of interest, namely the number of CD dimers *n*_CD_ or the number of CC dimers *n*_CC_ for pathways leading to *T* =4 or *T*=3. This choice allows us to distinguish off-pathway states; i.e. states with CC dimers in *T* =4 pathway and states with CD dimers in *T*=3 pathway. From the set of simulation trajectories at a given parameter set, we estimate the transition probability matrix **T**(*τ*), which has elements *T_ij_* that give the probability of a transition between a pair of states *i, j* in a lag time *τ* (see the Methods section).

With the constructed MSMs, we use transition path theory to identify dominant pathways, key structural intermediates, and points at which pathways are committed to certain morphology products as follows. The forward committor probability,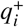 is the conditional probability that a trajectory that is in state *i* will visit a set of outcome states (denoted as B) before returning to the initial unassembled state. This is given by the solution to 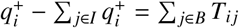, where *I* are all other states (the ‘intermediates’).

#### Commitor probabilities identify pathway selection hub states

Fig. 6 shows the committor probabilities for reaching the complete *T*=4 capsid *q*^+^ at two different dimer concentrations, *C*_tot_ = 10*μ*M (left) and *C*_tot_ = 20*μ*M (right). The most notable difference between the two concentrations involves the committor pobabilities of the states close to *T*=4 capsid (*n*_dimer_ > 100, *n*_CD_ > 25). While the majority of states end up in *T* =4 capsids (*q*^+^ ≈ 1) at the lower concentration, a fraction of these states have *q*^+^ ≈ 0 at higher dimer concentration; i.e. they remain trapped in other morphologies for the finite simulation time.

**FIG. 6.**
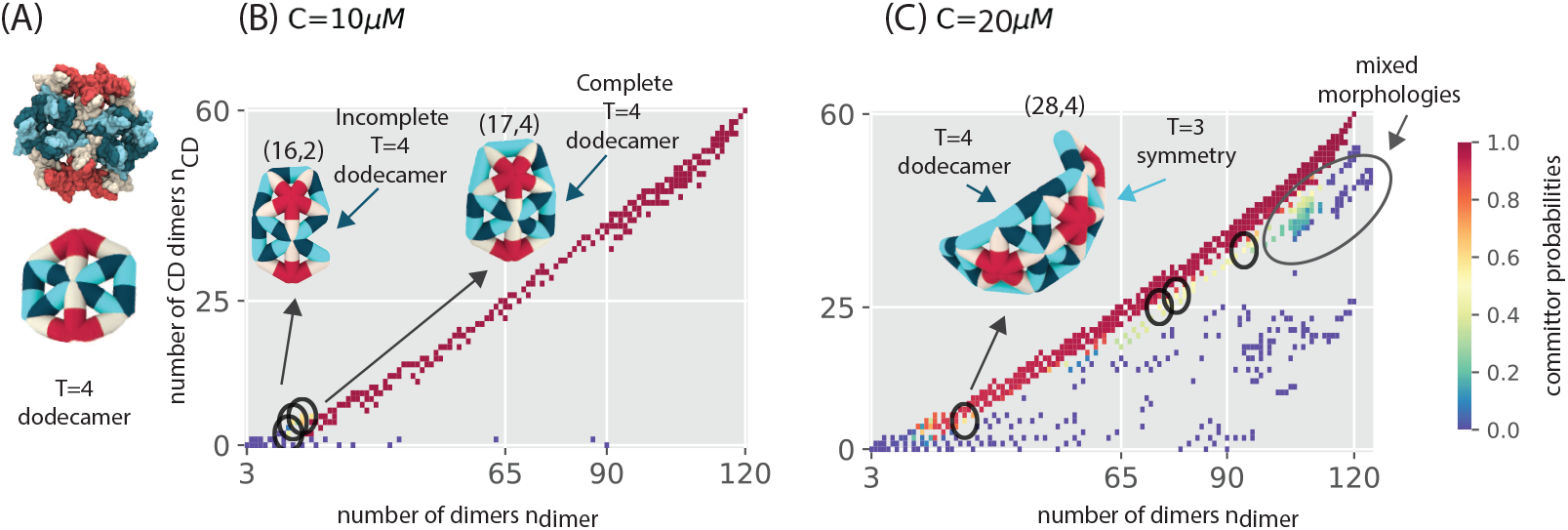
*T*=4 pathway ‘hubs’. **(A)** The structure of the ‘*T*=4 dodecamer’. **(B)** Committor probabilities for *T*=4 capsids *q*_*T*=4_ computed from Markov state model (MSM) analysis, shown as a function of the number of CD dimers *n*_CD_ and the total number of dimers *n*_dimer_ for *C*_tot_ = 10*μ*M at 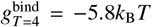 and 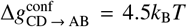. The committor probability =4 corresponds to the probability that a trajectory initiated at a given structure will form a complete *T*=4 shell before disassembling. Circles indicate the *T*=4 pathway ‘hubs’ (*q*_*T*=4_ ≈ 0.5), from which a trajectory is equally likely to form a *T*=4 capsid or other product morphology. Other snapshots are labeled as (*n*_dimer_, *n*_CD_). The (17, 4) intermediate in (B) is the smallest structure that has a complete *T*=4 dodecamer. **(C)** *T*=4 Committor probabilities for *C*_tot_ = 20*μ*M at similar parameters as in (B). At *C*_tot_ = 10*μ*M (B), the smallest hub states lack a complete *T*=4 dodecamer, while at the higher concentration (C) the smallest hub state has a mixed-morphology that includes a *T*=4 dodecamer. The region indicated by an oval at large sizes in (C) shows that at high concentrations there is a high propensity to form long-lived mixed-morphology structures with sizes between 85-140 dimers.

We identify the intermediates at which assembly pathways are most likely to diverge from formation of a *T* =4 capsid, resulting in an alternative assembly product, by identifying *hub states* as those for which the *T* =4 committor probability 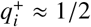. The five of these states with the highest forward flux (discussed next), as well as the smallest hub state, are shown with circles on the plots in Fig. 6(B-C). At the lower concentration, pathways diverging at the hub states typically result in *T*=3 capsids, whereas at the higher concentration assembly pathways are significantly more likely to diverge from *T*=4 to mixed-morphology states.

Comparison of the committor probabilities and assembly products at the different concentrations reveals that formation of a ‘*T* =4 dodecamer’ (snapshot in Fig. 6(A)) is the key event that determines whether pathways are more likely to proceed to *T*=4 or *T*=3 products. The *T*=4 dodecamer is the smallest relatively stable partial capsid intermediate that occurs in *T*=4 capsids but not *T*=3 capsids. The hub states at *C* = 10*μ*M (Fig. 6(B)) that lead to *T*=3 capsids occur before formation of a structure with a complete *T*=4 dodecamer, whereas hub states at *C* = 20*μ*M (e.g. the 28-dimer structure shown in Fig. 6(C)) occur after formation of a complete *T*=4 dodecamer, and consequently diverge to mixed-morphology malformed states. We find that with increasing concentration, pathways typically do not commit to the *T* =4 morphology until larger sizes and the size of the smallest hub state increases. Thus, increasing concentration results in higher probability of forming mixed-morphology structures across a range of large sizes (85-140 dimers). The increasing proportion of long-lived intermediates in this size range with increasing concentration is consistent with CDMS measurements [31].

The *T*=4 dodecamer was identified as a frequently observed intermediate along *T*=4 assembly pathways in AFM measurements of HBV dimers assembling on a mica plate [33]. The significance of this structure to formation of *T*=4 capsids in our simulations qualitatively agrees with this observation. However, unlike the AFM experiments, this intermediate evolves from a 10-dimer pentamer in our simulations. This difference may arise because the 10-dimer pentamer is a curved geometry which would be disfavored by the flat mica substrate used in the AFM experiments.

#### Forward flux reveals most probable assembly pathways

While the committor probabilities in Fig. 6(B-C) identify the ‘hub’ states for *T*=4 assembly, the *forward flux* 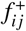 shows the relative probability of different reaction channels and hence the relative importance of different classes of assembly pathways. Fig. 7(A-B) shows the forward flux toward *T*=4 and *T*=3 morphologies respectively. The first order parameter is the number of capsid dimers. The second order parameter in each plot is chosen to show deviations from the target stucture pathway; i.e., the y-axis is number of CC dimers in a given state for *T*=4 pathways (Fig. 7(A)) and number of CD dimers for *T*=3 pathways in (Fig. 7(B)).

**FIG. 7.**
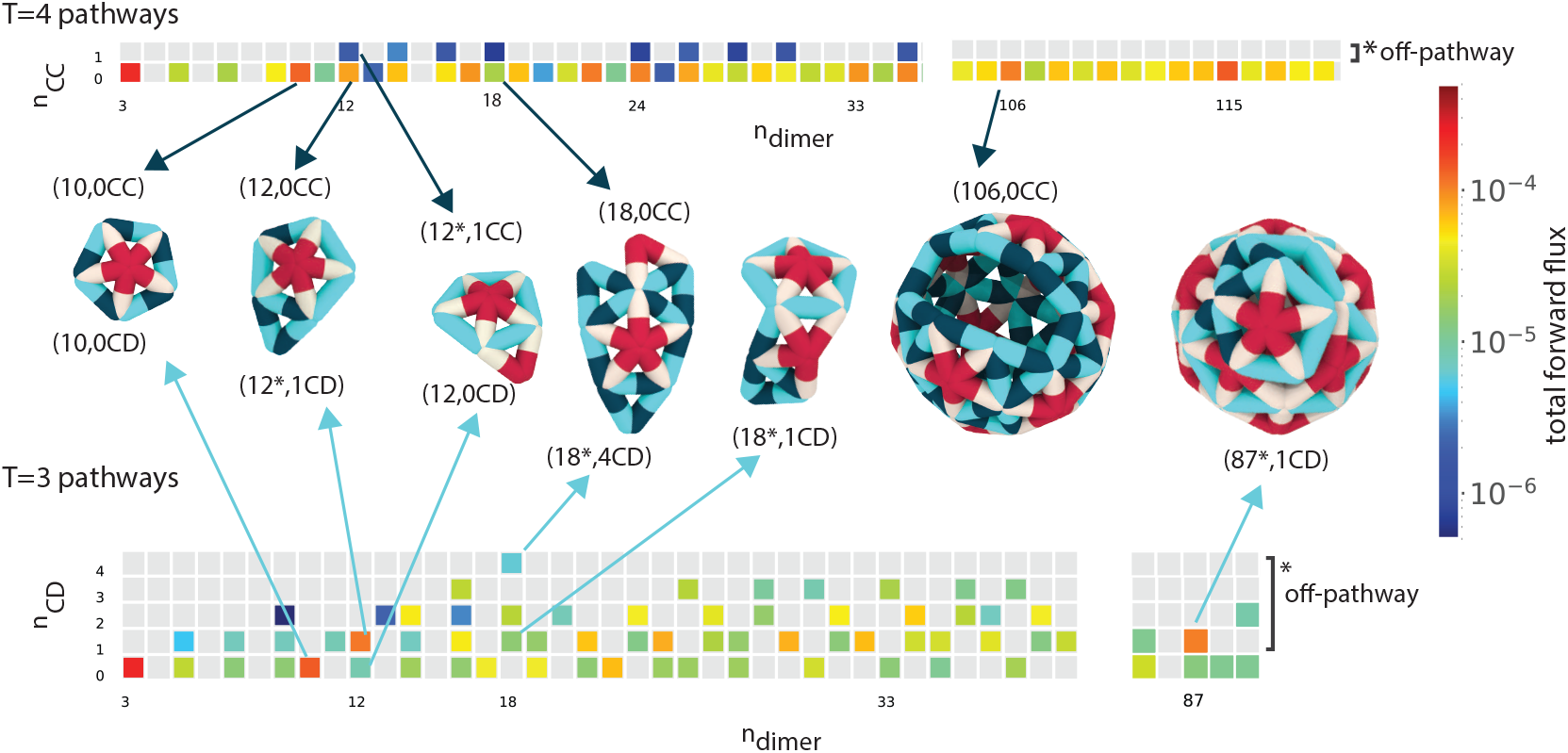
MSM analysis of intermediate stabilities and assembly pathway probabilities. Total forward flux to *T*=4 (top) and *T*=3 (bottom) capsids as the function of the number of dimers *n*_dimer_, and the number of CC (*n*_CC_) or CD (*n*_CD_) dimers for *T*=4 or *T*=3 pathways respectively. States with CC dimers (*n*_CC_ > 0) are off-pathway for *T*=4 assembly, and states with CD dimers (*n*_CD_ > 0) are off-pathway for *T*=3 assembly. For the *T*=4 pathway all the states with high forward flux are on-pathway states, while on *T*=3 pathways many of the states with high forward flux are off-pathway, and there are off-pathway states with up to 5 CD dimers in structures with high forward flux to a *T*=3 capsid. Simulation parameters are 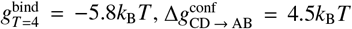, and C=20 *μ*M. Examples of assembly intermediates in the initial and final stages of the assembly are shown. Middle states are not shown for the clarity.

States with the largest flux between the initial and final configurations identify the most probable assembly pathways for each morphology. Interestingly, these states coincide with the long-lived intermediates discussed above (Fig. 5). The states with high forward flux toward *T*=4 capsids (Fig. 7(A)) can be understood by visualizing individual trajectories (movie S2). Typically, growth from one of these states begins with multiple off-pathway excursions, in which dimers bind with configurations incompatible with the final morphology, followed by stalling of growth, dissociation, and a return to the long-lived intermediate state, before eventual on-pathway growth.

While the states with highest forward flux on *T*=4 pathways are all on-pathway states (with no CC dimers), the *T*=3 pathway exhibits several off-pathway states with high forward flux. Interestingly, the 18-dimer (18*,4CD) state on the *T*=3 pathway is actually an on-pathway *T*=4 intermediate, but it can also generate pathways that form mixed-morphologies that undergo annealing to form *T*=3 capsids. Another example of such a trajectory is shown in Fig. 5(B) and movie S5) where the local *T*=4 symmetry remains in the structure until the final steps of assembly. This pathway includes a long-lived intermediate that has 90 dimers but is incomplete due to the presence of local *T*=4 regions. In contrast, we do not observe any trajectories that transition from a *T*=3 pathway to a complete *T*=4 capsid.

## DISCUSSION

We have investigated the assembly pathways of HBV capsid proteins using a coarse-grained model, informed by atomic-resolution data from molecular dynamics simulations and structure-based estimates of binding affinities for different protein conformations. The simulations reproduce key assembly products observed in experiments, including polymorphic assembly into *T*=3 and *T*=4 icosahedral structures and non-icosahedral complexes. Notably, our simulations predict the structural characteristics of the off-pathway intermediates, which could not be inferred from the CDMS experiments [29–32]. At the optimized values of the elastic moduli and binding energies, the off-pathway products have a mixed-morphology comprising a combination of local *T*=4 and *T*=3 symmetry environments. The curvature of such structures is not geometrically compatible with self-closure, leading to shells with defects or holes, as well as overgrown structures in which the assembling structure spirals around itself until assembly is stalled due to excluded volume and elastic strain. These observations elucidate the experimental finding that metastable asymmetric intermediates can convert into icosahedral capsids over several days [31], as well as slow capsid closure during the final stages of HBV assembly observed using SAXS experiments [36]. The simulations also predict key factors in the assembly of large aberrant structures with lower curvature observed in HBV experiments with CpAM modulators, and demonstrate that the combination of specific molecular scale interactions and flexibility, in particular the relatively small bending modulus in comparison to other virus capsids, enable HBV to undergo high fidelity assembly into the infectious capsid structure while retaining the capability to form other polymorphs.

The simulations suggest that due to this flexibility, a coupling between the protein conformational state and its protein-protein interactions is critical for determining assembly pathways and products. These results suggest potential new interpretations of existing experimental data. For example, existing models have not been able to explain the experimental observations that higher proportions of *T*=3 capsids assemble at higher ionic strengths, and in ammonium acetate buffer compared to NaCl [36, 137].

The observation from our simulations that 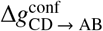 has a strong effect on morphology selectivity suggests that the ionic strength shifts the conformational equilibrium toward CD dimer conformations. That is, increasing the salt concentration decreases 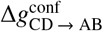, and thereby increases the driving force for fivefold dimer coordination during assembly process, thus favoring pathways that lead to the *T*=3 morphology. These results are also consistent with recent suggestions that coupling between conformational interconversion and interaction strengths provides an important means of regulatory control over the timing and robustness of assembly [112, 134, 140, 141]. Moreover, Biela et al. [142] recently demonstrated that assembly of larger capsids from MS2 capsids (*T*=4 and D5) could be achieved by engineering insertions into the capsid protein that likely shift the capsid protein conformational equilibrium (analogous to changing our parameter 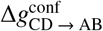).

The need for conformational specificity when accounting for HBV material properties sheds new light on previous results for a model of icosahedral capsid assembly with only one subunit species, which found that much higher values of the bending modulus, corresponding to much smaller values of the Foppl von Karman number (FvK < 0.25, compared to FvK ≈ 500 for HBV), were required to observe assembly into *T* =4 structures [91, 143]. Here we find that *T*=4 capsids assemble with high yields at FvK ≈ 500, provided that the dimer subunits can adopt quasi-equivalent conformations with different binding affinities.

Trajectory analysis shows that, while the products of HBV assembly can be qualitatively classified into *T*=4 capsids, *T*=3 capsids, and long-lived malformed structures, the assembly pathways all have key features in common. We identified hub states, or intermediates at which assembly pathways frequently diverge from *T*=4 pathways, as well as structural features of such hub states that determine the classes of products they will form. A key feature is whether a hub state intermediate contains a dodecamer of dimers in the quasi-sixfold arrangement found in *T*=4 capsids (the *T*=4 dodecamer in Fig. 6(A)).

Trajectories that form such an intermediate will go on to form either *T*=4 capsids or malformed structures, whereas trajectories that diverge from the *T*=4 pathway before forming a complete *T*=4 dodecamer form either *T*=3 capsids or malformed structures. This prediction is consistent with recent AFM observations of HBV capsid protein assembly [33]. Knowledge of hub states could suggest new antiviral drug targets.

### Outlook

The simulation results and buried surface area estimates suggest that, although the different quasi-equivalent conformations of HBV capsid proteins have high structural similarity, their differences play a key role in guiding assembly pathways. This conclusion is further supported by the fact that the antiviral assembly effectors (CpAMs) bind preferentially to interfaces between certain conformational pairs (e.g. within the quasi-sixfold interfaces). It would thus be of great interest to extend recent atomic-resolution simulations of CpAM-HBV capsid protein interactions [144, 145] to investigate the molecular-scale factors that couple the protein conformational state to its interactions. This information could facilitate designing more effective assembly effectors.

The importance of conformational heterogeneity in HBV assembly highlights the inherent trade-off between minimizing complexity of a self-assembly reaction and maximizing selectivity for target structures. Recent experiments and simulations of synthetic subunits designed to assemble into capsids and tubules demonstrated that multiple species with species-specific subunit-subunit interactions can significantly increase specificity for a target geometry, by blocking assembly pathways that would lead to other geometries with similar thermodynamic stabilities [146–148]. However, the additional information content associated with encoding for multiple species or conformations incurs extra costs, such as material and design costs in the synthetic realm, or additional selective presssure on protein sequences in natural systems. Understanding how this trade-off has shaped evolution in other natural systems would provide important information for developing treatments for pathogenic diseases that work by redirecting assembly pathways, and could guide more efficient design of synthetic self-assembly systems.

## METHODS

### Coarse grained (CG) model

In our CG model, an assemblage is represented as a triangulated elastic sheet, similar to models previously used for assembly of microcompartments [122], viral capsids [92], and geometrically frustrated assembly [124]. However, while these previous models were based on association and dissociation of triangular subunits, the basic assembly unit in our model is an HBV dimer, which corresponds to an edge in the elastic network. This choice is based on the fact that dimers are the basic assembly unit in HBV capsid assembly [120, 121].

In particular, each HBV dimer is modeled as an edge with two asymmetric half-edges. Half-edges have orientations (e.g. CD conformation edges comprise CD and DC half-edges in the model), and the edge-edge bindings are type- and order-dependent (e.g. 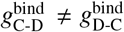). This is an important feature for modeling HBV assembly, because the dimer-dimer interfaces in HBV capsids are asymmetric — one dimer is ‘capped’ at one end by the other dimer. The model *T* =4 capsid has 120 edges corresponding to the 120 dimers in a complete *T* =4 HBV capsid.

The energy for a model capsid configuration is given by:

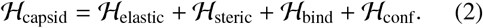

The elastic energy accounts for the harmonic potentials for edge length fluctuations *l*, dihedral angles *ϕ* between the planes of each pair of adjacent triangles, and binding angles *θ* between each pair of edges meeting at a vertex, with moduli *κ_l_, κ_θ_* and *κ_ϕ_* respectively:

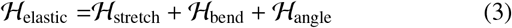

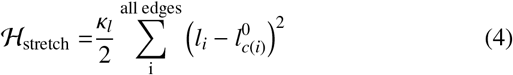

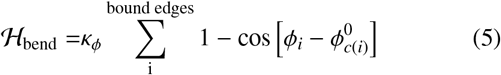

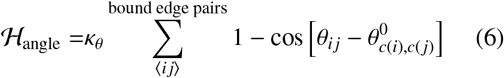

with *c*(*i*) the dimer conformation for edge index *i* and *c*(*i*), *c*′(*j*) the conformations of two bound edges, with *i, j* the edge indices. Minimum energy values for edge lengths, dihedral angles, and binding angles are given by 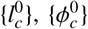, and 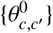, with *c* ∈ {AB, CD} the set of distinct dimer conformations (as discussed below, the geometric parameters are the same for CC and CD conformations). The associated moduli, *κ*_l_ = 4200 *k_B_T*/*σ*, *κ_ϕ_* = 40 *k*_B_*T*, and *κ_θ_* = 800 *k*_B_*T* are set to be independent of edge conformation (See SI Model section and Fig. S3 for parameter values of equilibrium lengths and angles).

The term 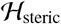 represents the excluded volume interaction of dimers, and is represented by hard-sphere excluders with positions relative to the edge axis that are based on the dimer structure (see SI Fig. S1). Excluders move as a rigid body with the edge position and orientation. Configurations in which excluders overlap have infinite energy and thus are forbidden.

The binding energy 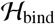 is the sum of binding energies between pairs of bound edges (*i, j*), with a binding affinity that depends on the edge conformations, *g*_*c*(*i*′),*c*′(*j*)_. The half-edge data structure used in our implementation has an orientation and thus efficiently represents the conformation- and order-dependant binding energies in HBV capsids.

The term 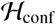 represents intra-dimer conformational free energy landscape; i.e., the relative equilibrium populations of different conformational states for the capsid protein dimers. We specify this distribution according to the equilibrium constant and corresponding free energy difference between pairs of conformational states, with CD as the zero free energy reference conformational state. For example, for the two dimer conformations found in the *T*=4 capsid, AB and CD, we define 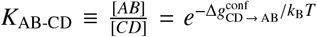, where *K*_AB-CD_ is the equilibrium constant for interconversion between the AB and CD conformations and 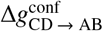 is the corresponding free energy difference between the two conformations. We set the free energy of the CC conformation (found in *T*=3 capsids) equal to that of CD, i.e. *K*_CC-CD_ ≡ [CC]/[CD] = 1, based on the high degree of structural similarity between CD and CC conformations and to reduce the number of model parameters. Thus, the total conformational energy of an assembly is given by the number of AB dimers *n*_AB_ in the structure, 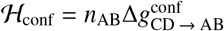.

### Monte Carlo simulations

To model the limit of dilute, noninteracting capsid structures typical of productive capsid assembly reactions [58, 126], we perform dynamical grand canonical Monte Carlo (MC) simulations of a single assembling shell in exchange with free dimers at fixed chemical potential *μ*, which sets the bulk dimer concentration [122]. The initial configuration for each simulation is three edges (dimers) bound in a triangle geometry, with the initial conformation of each triangle chosen randomly. The MC algorithm includes 7 moves, which include association or dissociation of either single dimers or pairs of dimers, binding or unbinding of the dimers in the shell, conformational switches of individual dimers, and thermal relaxation of the shell by vertex displacement moves (SI Fig. S2). Moves are accepted or rejected according to the Metropolis-Hastings acceptance criteria [149, 150]:

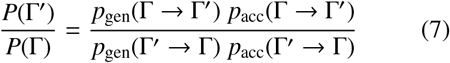

where Γ and Γ′ denote the initial and trial states, the probability of a state with *n*_dimer_ edges is 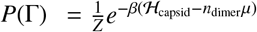 with *Z* the grand canonical partition function, and pgen and pacc are respectively the probabilities for generating and accepting trial moves.

### Estimating CG model parameters from an AA simulation of a complete HBV capsid

We estimate the parameters for 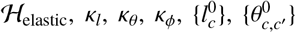, and 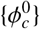, by comparison against AA molecular dynamics simulations of a complete *T*=4 HBV capsid [1]. We coarse-grain the data from the AA simulations in two steps. First, we select the C-*α* atom of residue 132 of each capsid protein monomer. The rationale for using residue 132 as an anchor residue is two-fold. One, it provides a convenient point of reference due to its position near opposite ends of the dimer’s principle axis. Two, it is a well-studied residue; amino acid substitutions at this site have shown measurable effects on capsid assembly and capsid stability [151]. These 240 points are clustered based on proximity using the scikit DBSCAN algorithm [2], which identifies the fivefold and quasi-sixfold axes. The center-of-mass of each cluster is then assigned to a vertex in the CG capsid. Calculating the set of CG edge lengths, dihedral angles, and binding angles for each residue based on 50,000 conformers from the 1-*μ*s AA simulation provides the data for estimating the equilibrium distribution of these parameters in the CG model (see Fig.1(C) and movie S1).

We perform MC simulations on complete *T*=4 capsids to estimate the corresponding distribution of edge lengths and angles in the CG model. In particular, we use the complete *T*=4 capsid mapped from a frame in the AA simulation as the initial condition, and only perform vertex displacement moves to relax the structure. In each simulation, we initially perform 50,000 MC sweeps to equilibrate the system and then an additional 50,000 sweeps to estimate the edge length and angle distributions. We optimize parameter values by minimizing the Kullback–Leibler divergence of the CG and AA distributions for each quantity, using the Tree-structured Parzen Estimator (TPE) algorithm in Hyperopt package[152].

As shown in Fig. S3 the distribution of edge lengths is bimodal, and thus the edge-length distribution can only be fit by a CG model with at least two edge types (corresponding to two different conformations, AB and CD). Thus, in addition to the elastic moduli, we fit equilibrium edge lengths for both conformations 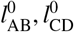, dihedral angles 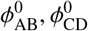, and binding angles 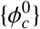 with (*c, c′*) ∈ {(BA, AB), (AB, CD), (CD, BA), (DC, DC)}.

In addition, to the binding angle conformation pairs listed above for *T*=4 and *T*=3 capsids, structures of HBV dimers in drug-mediated assembled structures (and similarly hexagonal sheets of HBV dimers) include a binding angle pair CD, CD, for which we set 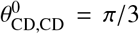 since the sheet is composed of equilateral triangles.

### Estimating binding energy parameter values

Atomic-resolution structures of different HBV capsids and assemblies [23, 125] show that there are significant structural differences between the binding interfaces for different dimer conformations. Therefore, in our model the dimer-dimer binding affinities depend on the dimer conformations. Since the dimers are not head-tail symmetric, the binding affinities also depend on the relative orientations of the monomers within each dimer. Thus, there are 16 possible binding arrangements for two dimer types AB and CD. However, only seven of these binding arrangements can be found in available structures of HBV capsids: (BA-AB, AB-CD, CD-BA in both *T* =4 and *T*=3 capsids; DC-DC in *T* =4 capsids; AB-CC and CC-BA in *T*=3 capsids, and CD(CC)-CD(CC) in drug-mediated assembled structures (and similarly hexagonal sheets of HBV dimers) [23, 125, 131, 132].

To estimate the relative difference in binding affinity between different dimer-dimer conformation pairs, we use the buried surface area computed from atomic-resolution structures using PDBePISA, a tool for examining macromolecular interfaces [153, 154]. Table I shows relative binding energies calculated from two different *T* =4 HBV capsid crystal structures 2G33[23] and 6UI7[125], where we have set the reference binding energy parameter 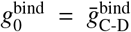. In *T*=3 capsids, AB-CC and CC-BA have similar buried surface areas to AB-CD and CD-BA in *T* =4 capsids [125] (Table S2), and thus we consider them to be the same in the model. The CD-CD interaction is not present in *T*=4 or *T*=3 capsids, but calculation of the CD-CD binding affinity for CD conformations in the *T*=4 structure [23] gives 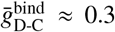. We have optimized this parameter in Fig. 2 (A) and (B) and set the 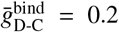. Similarly, estimates for all other binding affinities result in small values 0.01-0.1 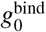; thus, to limit the number of model parameters we set all other interactions to 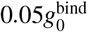.

### Mapping the free model parameters to experimental values

The structure and dynamics-based procedures described thus far to set model parameters leave two free parameters, 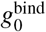, which controls the mean inter-dimer binding affinity, and 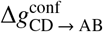 the intra-dimer conformational interconversion free energy. These parameters together set the mean free energy of a dimer in a capsid ground state configuration (meaning subunits are at their equilibrium position so that 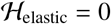), according to:

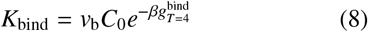

with

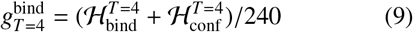

where 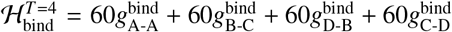 and 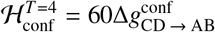, *v*_b_ is the binding volume parameter (see SI), and *c*_0_ = 1M is the standard state volume. *c*_0_ = 1 M is also used for calculating the chemical potential *μ* = ln(*c*/*c*_0_).

The mean dimer-dimer binding free energy corresponding to *K*_bind_ has been estimated from experimental measurement of the equilibrium capsid assembly yields as a function of total subunit concentration under different experimental conditions (e.g. [120]). However, the assembly equilibrium depends on both 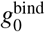 and 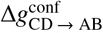, and it is likely that both of these parameters depend on solution conditions [120, 133, 134, 140, 155]. Thus, it is not straightforward to estimate both of these parameters from the experimental data. Instead, we consider 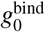 and 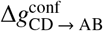, but note that their ranges and relative values are constrained by experimental data according to Eq. (8).

### Markov State Models (MSMs)

To build MSMs, we start by partitioning configurations from the simulation trajectories into states, such that configurations that interconvert rapidly are collected into the same state. This separation of timescales ensures that the model is roughly Markovian, meaning that the probability of transitioning to a new state only depends on the current state, on timescales longer than a ‘lag time’ *τ* that corresponds to the relaxation timescale within a state.

We cluster configurations based on geometric criteria and the different types of dimers in the assemblage and use two order parameters to perform state decomposition. The first order parameter is the number of dimers *n*_dimer_ in a partial capsid structure. For the second order parameter, we use the number of CD dimers *n*_CD_ and the number of CC dimers *n*_CC_ and construct different MSMs for the analysis of *T*=4 and *T*=3 pathways.

We then calculate the transition matrix **T**(*τ*) from the ensemble of MC trajectories as follows. We compute the *count matrix* **C**(*τ*), where each element *C_ij_* is the total number of transitions from state *i* to state *j* measured at a lag time *τ*. The *transition matrix* **T**(*τ*) is then calculated from by column-normalizing the count matrix **C**(*τ*). The time-dependent state probabilities can then be calculated using spectral decomposition of the transition matrix according to:

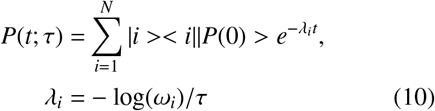

where *ω_i_* is the *i*-th eigenvalue of **T**(*τ*), < *i*| and |*i* > are the corresponding left/right eigenvectors, and the implied timescale, 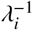 corresponds to the relaxation timescale for eigenmode *i*.

Calculating committor probabilities for particular structures (e.g. the complete *T*=4 capsid) is complicated by the fact that there are multiple absorbing states, since closed shells and some other mixed-morphology states will not disassemble on timescales accessible to any of our simulations. To prevent degenerate eigenvectors, we add the initial state at the end of all trajectories ending in absorbing states other than the target structure of interest. That is, when calculating 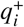 for *T*=4 structures we add the initial state at the end of any trajectory which concludes in an absorbing state (including *T*=3 structures); similarly when computing 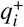 for *T*=3 we add the initial state to the end of trajectories that end in absorbing states including *T*=4 structures.

## ACKNOWLEDGMENTS

This work was supported by Award Number R01GM108021 from the National Institute Of General Medical Sciences and the Brandeis Center for Bioinspired Soft Materials, an NSF MRSEC, DMR-2011846. Computational resources were provided by NSF XSEDE computing resources (XStream, Bridges, and Comet) and the Brandeis HPCC which is partially supported by DMR-2011846. J.A.H.-P. acknowledges funding from the National Institute Of General Medical Sciences under Award Number P20-GM-104316.

## SUPPORTING INFORMATION

### Implementation of the coarse-grained (CG) model

We implement our model of dimer subunits using the half-edge data structure (HE) [156], which is a doubly connected edge list (DCEL)[157]. Each edge in the model corresponds to a protein dimer, and each edge consists of two half-edges. We consider two types of edges: The type AB edge consists of AB and BA half-edges in opposite directions and represents the AB dimer conformation in *T*=3 and *T* =4 capsids; the type CD edge consists of CD and DC half-edges in opposite directions and represents the CD dimer in *T*=4 capsids or the CC dimer in *T*=3 capsids (Figure S1(A-B)). Each of these edge types is associated with an equilibrium length, 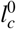 with *c* = AB, CD labeling the conformation, and each interior edge (located between two triangle faces in the structure) has an equilibrium dihedral angle 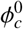. As noted in the main text, due to the high structural similarity between CD and CC dimers, we implement them with the same values of 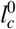 and 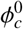 and thus associate them with a single edge type. Throughout the article a CD/CC dimer is called a CD dimer except when it has a (CC/DC)-BA or AB-(CC/DC) interaction S1(C), as these two interactions are observed in *T*=3 capsids but not in *T* =4 capsids.

The advantage of the half-edge data structure, in comparison to related triangular sheet models [90, 122], is that the two half-edges allow different parameters for protein-protein interactions between different monomer conformations. In particular, with 4 halfedges *c* ∈ {AB, BA, CD, DC}, there are up to 16 different values for binding affinities *g_cc′_* and equilibrium binding angles *θ_c,c′_*, for dimer-dimer interactions. Note that the edges are asymmetric, and thus *g_cc′_* and *θ_cc′_* depend on the order of the two half-edges involved; i.e., in general *g_c,c′_* ≠ *g_c′,c_*.

Now we explain how the combination of 2 edge types and different interaction types are sufficient to represent the range of HBV protein conformations observed in different assembly morphologies (*T*=3, *T* =4, and the sheets, tubules, and other aberrant structures observed in HBV-CpAM assemblies).

Table S1 shows the binding affinity matrix for the possible half-edge-interactions in the model. Each table entry shows the binding affinity, relative to the D-C binding affinity, for a given half-edge and the half-edge in the nearby dimer that it interacts with. The ‘Structure’ column gives the atomic-resolution structure from which we estimated the buried surface area for the calculation of the binding affinity. For interactions that involve CC dimers in *T*=3 capsids, CC dimers have similar buried surface area as the CD dimer in *T* =4 capsids in the corresponding local geometry; i.e., CC-BA and AB-CC interfaces in *T*=3 capsids have very similar buried surface areas as CD-BA and AB-DC interfaces in *T*=4 capsids (see Table S2 and reference [125]).

**TABLE S1.** Interaction angles and the relative binding affinities for the different conformations of interacting dimer pairs observed in assembled structures.

| c(h) | c'(next-h) | contact | $g_{cc'}/g_0^{\text{bind}}$ | $\theta_{c,c'}^0$ | structure |
| --- | --- | --- | --- | --- | --- |
| BA | AB | A-A | 1.3 | 1.17 | T4,T3 |
| AB | CD(CC) | B-C | 0.9 | 0.98 | T4,T3 |
| CD(CC) | BA | D-B | 1.1 | 0.98 | T4,T3 |
| AB | DC(CC) | B-D | 0.9 | 0.98 | T3 |
| DC(CC) | BA | C-B | 1.1 | 0.98 | T3 |
| DC | DC | C-D | 1 | 1.05 | T4 |
| CD | CD | D-C | 0.2 | 1.05 | w/drug |
| other | other |  | 0.05 | 1.05 | w/drug |

**TABLE S2.** Comparison of dimer-dimer binding affinities in *T*=4 and *T*=3 capsids.

| | $\bar{g}_{A-A}^{\text{bind}}/k_B T$ | $\bar{g}_{D-B}^{\text{bind}}/k_B T$ | $\bar{g}_{B-C}^{\text{bind}}/k_B T$ |
| --- | --- | --- | --- |
| 6UI7 $T=4$ capsid [125] | -8.36 | -7.35 | -6.11 |
| | $\bar{g}_{A-A}^{\text{bind}}/k_B T$ | $\bar{g}_{C-B}^{\text{bind}}/k_B T$ | $\bar{g}_{B-C}^{\text{bind}}/k_B T$ |
| 6UI6 $T=3$ capsid [125] | -8.42 | -7.71 | -6.52 |

The equilibrium interaction angles 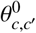 for dimerdimer interactions observed in *T*=4 and *T*=3 structures are optimized from all-atom (AA) simulations of *T*=4 capsid as explained in the main text. All other equilibrium interaction angles are set to *π*/3 since they generally occur in flat or nearly flat hexagonal structures.

### Monte Carlo (MC) Simulations

Our grand canonical Monte-Carlo simulation implementation is adapted from Refs. [122, 124], in which the triangular sheet is represented by edges and vertices. In our model, each edge is associated with two vertices, which together have 2 × 3 degrees of freedom. Any two bound edges in the shell share a vertex, and thus together have 3 × 3 degrees of freedom. For a shell with *n*_dimer_ edges, the grand canonical probability density is

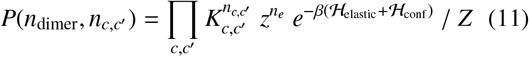

where *c* ∈ {AB,BA,CD,DC} are the different half-edge types (representing their conformations and directions) and *n_c,c′_* is the number of interactions within the shell configuration between pairs of dimers with *c* and *c*′ conformations respectively. 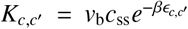, with *ϵ_c,c′_* the binding affinity of interacting half-edges with conformations *c* and *c′, v_b_* the binding volume and *c*_ss_ the standard state concentration. *z* = *e*^-*βμ*^/*λ*^6^ is the activity with the chemical potential *μ* = log(*c*_d_/*c*_ss_) where cd is the dimer concentration in solution. The standard state concentration is set to *c*_ss_ = 1*M*. The term 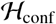 accounts for the conformational free energy difference of the two types of dimers in the shell, and 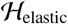 accounts for the elastic energy penalty for deviations from the ground state of the sheet.

**FIG. S1.**
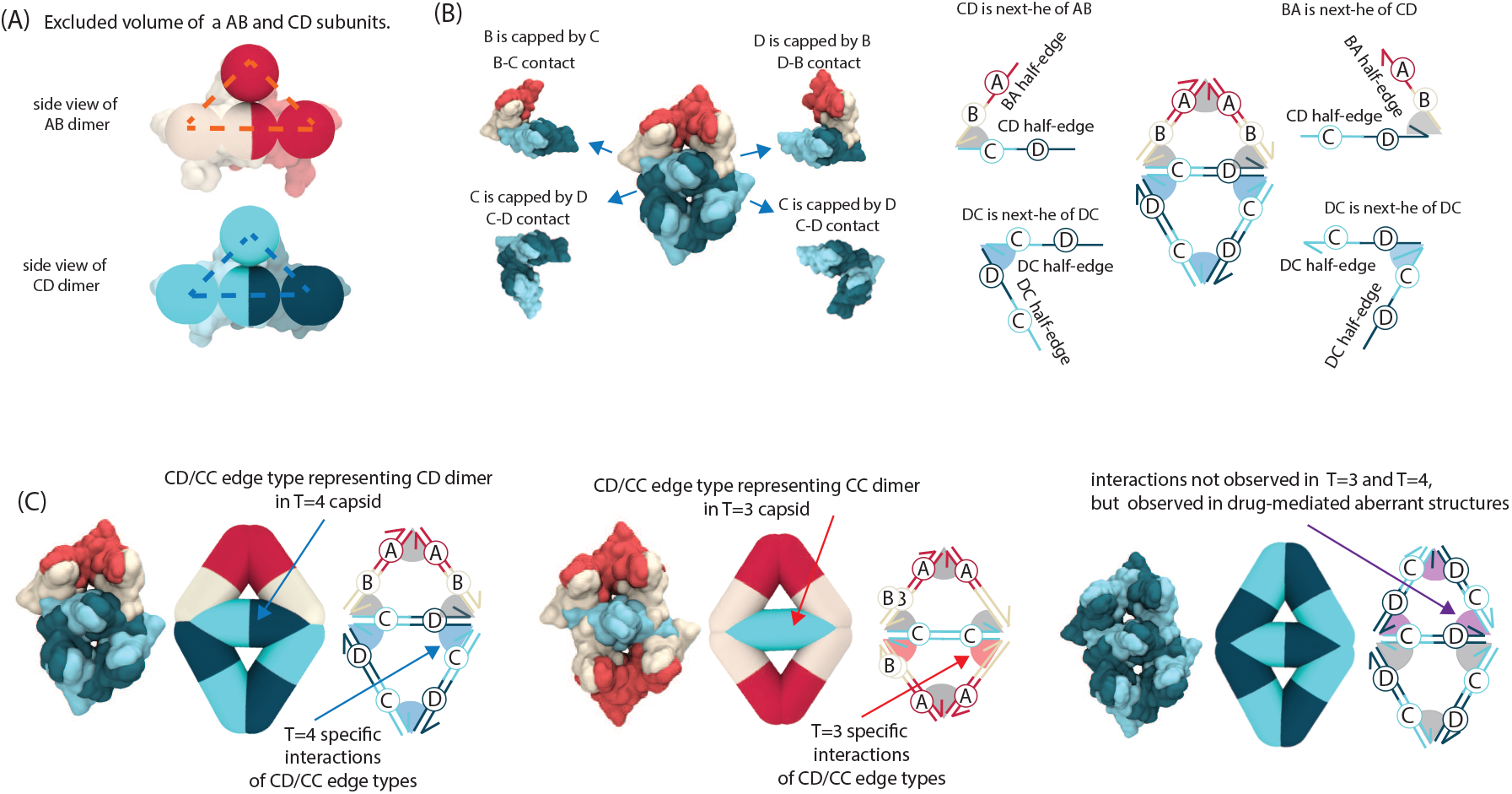
Mapping the half-edge data structure to HBV dimers. **(A)** Side view of overlay of HBV AB and CD dimers and model subunits with excluder pseudoatoms. Subunits are also prevented from overlapping with each other by forbidding the plane shown by dashed lines to intersect the corresponding planes on other subunits. **(B)** The four different contacts of the middle CD subunit in a T=4 intermediate structure (left) and the relevant implementation of each contact by the half-edge data structure (right). In an HBV capsid, each dimer is capped in two of its four contacts, in a specific order as shown in the figure, which is represented by the contacts of two oppositely directed half-edges within an edge. **(C)** The two edge types, AB and CD, are shown along with the different edge interactions that they can make, to represent different conformations and interactions of HBV capsid protein dimers that are observed in available structures: T=4 (left), T=3 (middle), and hexameric sheets and drug mediated structures (right).

The binding volume *v*_b_ is 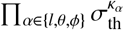 with 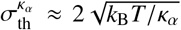the thermal length scales corresponding to elastic and bending moduli (*κ*_l_, *κ_θ_*, and *κ_ϕ_*) that govern edge geometry fluctuations.

In each MC step, a trial Monte-Carlo move *v* is chosen randomly from the list of Monte-Carlo moves (described below) according to its relative trial rate 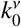. The trial moves are accepted/rejected based on the Metropolis-Hastings algorithm and detailed balance is ensured as presented in Eq. 7 of the main text.

### Parameter optimization from AA molecular dynamics trajectories

We set the values of 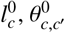 and 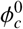 in our CG model from the mean values of the corresponding parameters in the CG representation of AA simulations of the *T*=4 HBV capsid. Parameter values of equilibrium lengths and dihedral angles are as follows: 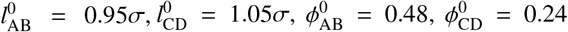, and all 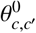 are listed in table S1. Other model parameters *κ*_l_, *κ_θ_, κ_ϕ_* cannot directly be inferred from the AA simulations. To optimize the model parameters, *κ*_l_, *κ_θ_* and *κ_ϕ_*, we perform simulations on a closed shell with a fixed topology corresponding to a *T* =4 icosahedral capsid. After equilibrating the shell, we optimize parameters by minimizing the difference between the distributions of thermal fluctuations in the CG model simulations and the AA molecular dynamics trajectories on a *T*=4 HBV structure. Because the topology is fixed in the CG simulations, we only perform vertex relaxation moves (described below). We perform *N*_eq_=50,000 sweeps during the equilibration phase of the simulation, where a sweep is defined as *n*_vert_ = 80 trial vertex displacements, so that each vertex on average will have undergone one trial move during a sweep. We measure the extent of equilibration by measuring the fluctuations of the elastic energy per dimer in the last 10,000 sweeps of the equilibration simulations 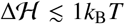 (Fig. S3(B). We then perform an additional *N*_CG_=50,000 sweeps to estimate the distribution of thermal fluctuations.

#### Optimization routine

Optimizing the CG model parameters is done by comparing the distributions of 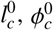, and 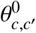 measured from AA molecular dynamics simulations and the MC simulations of complete *T* =4 capsids. Using an optimization method, and starting from a guess for the values of *κ*_l_, *κ_θ_*, and *κ_ϕ_*, the optimizer searches for the values that minimizes the loss function 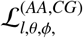. The loss function is the average Kullback–Leibler divergence *D*_KL_ of the two distributions of AA and CG simulations of each parameter multiplied by *λ* = 100 (to avoid numerical issues)

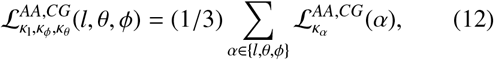

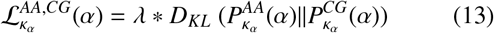

and 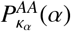 and 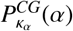 are distributions of lengths and angles of the CG and AA simulations respectively at a parameter set *κ*_l_, *κ_ϕ_* and *κ_θ_*. To explore the parameter space efficiently and to avoid becoming trapped in local minima, we use the Tree-structured Parzen Estimator (TPE) optimization algorithm of Hyperopt [152], which is a Bayesian optimization method. After 1000 trials, the loss function was minimized 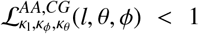 for 3000 < *κ*_1_/*σ k*_B_*T* < 5000, 300 < *κ_θ_*/*k*_B_*T* < 1500 and 20 < *κ_ϕ_*/*kt* < 50. Among the points with minimal loss function, we chose the optimal parameter set for performing simulations, where the three loss functions had similar values with the standard deviation 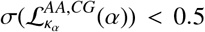. CG distributions for each quantity at optimal stretching, bending, and binding angle moduli *κ*_l_/*σ*^2^*k*_B_*T* = 4200, *κ_ϕ_*/*kt* = 40, and *κ_θ_/kt* = 800 are shown against the AA distributions in Fig. S3(C).

#### Comparison to a model with only subunit type

We also performed the parameter optimization for a model with a single type subunit in the capsid, similar to models used in [92, 122]. The CG distributions for 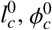, and 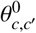 at optimal stretching and bending moduli *κ*_l_/*σ*^2^*k*_B_*T* = 2600 and *κ_θ_*/*k*_B_*T* ≈ 150, are shown against the AA distribution in Fig. S3(D). The minimum value of the loss function in the one-subunit type model is an order of magnitude larger than in our two-edge type model. Moreover, at these parameters, formation of fivefold vertices is unfavorable and assembly of empty *T*=4 capsids was unsuccessful, as also reported in [92].

### Shell assembly simulations

To simulate capsid assembly dynamics, we intersperse a variety of MC moves, described in the next section, that are designed to capture the physical dynamics of assembling subunits. Each simulation begins with an initial state comprising three edges in a triangular face. This state is known to be a highly populated intermediate, and is estimated to be the critical nucleus for HBV capsid assembly [33, 119, 137]. Each simulation contains a single assembly, which undergoes exchange of subunits with a reservoir according to the grand canonical probability density, Eq. (11). The simulations are performed for a maximum of 2 × 10^8^ sweeps, where a sweep is defined as a set of trial moves consisting of addition/removal, binding/unbinding, shell relaxation, and conformational switch moves such that each edge on average will have undergone one conformational switch move and each vertex on average will have undergone one vertex displacement move. Simulations are stopped early if the assembly forms a closed structure, defined as a structure in which every edge has its maximum number of four interactions. Simulations are also stopped early if the assembly becomes stalled in a sufficiently long-lived intermediate, defined as a structure with *n*_dimer_ ≥ 90 edges for which no growth has occurred for at least *n*_stall_ = 20/*α* sweeps where *α* = (*n*_dimer_ - 35)/*n*_sweep_ is the average growth rate in a given simulation after the structure reached *n*_dimer_ = 35 edges.

**FIG. S2.**
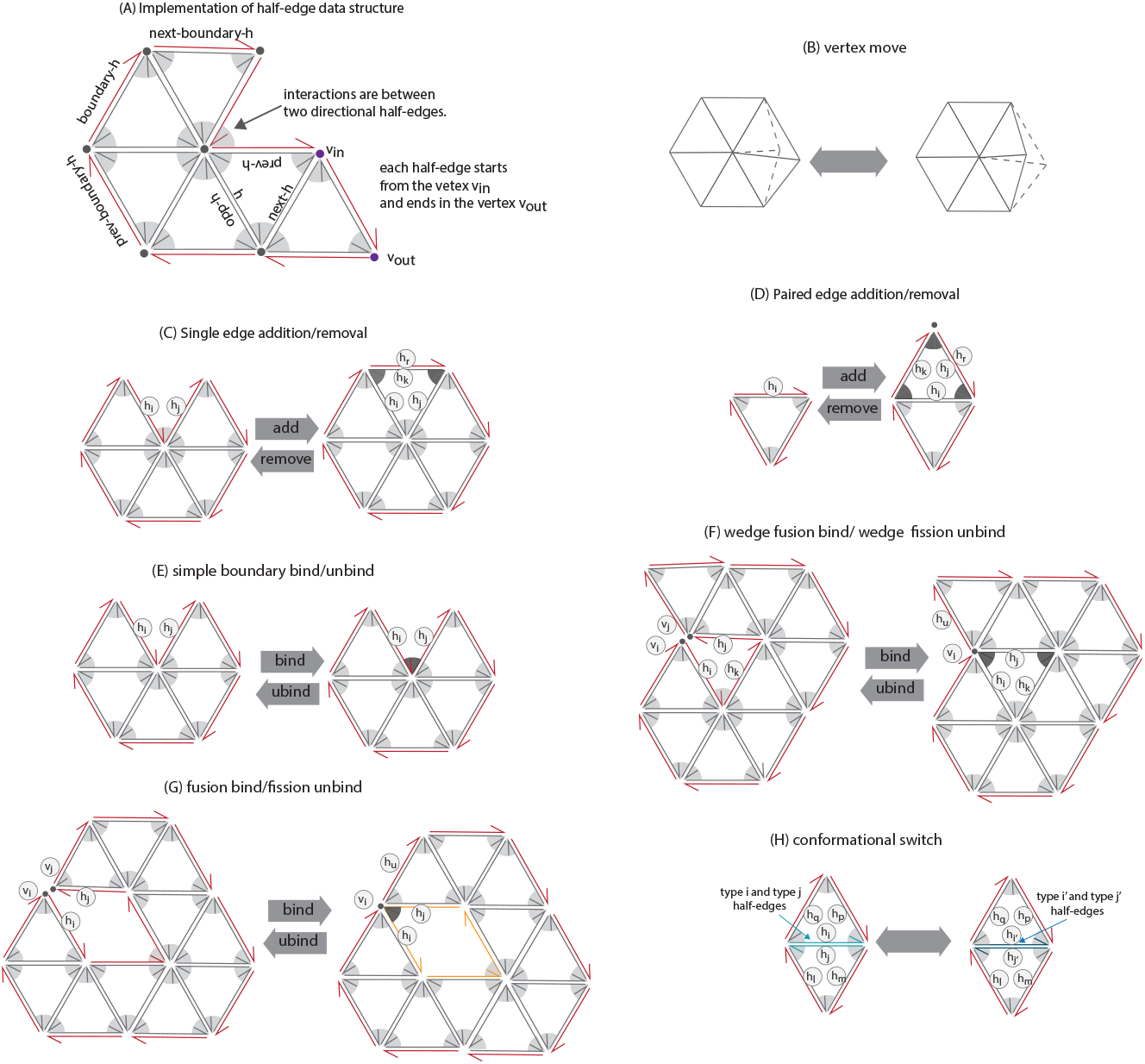
Schematic of the half-edge implementation and the MC moves. **(A)** Half-edge implementation. Each edge in the model is composed of two half-edges. Half-edges with no open ends are designated as *non-boundary half-edges*, and drawn as grey arrows. For a compact notation, for a given half-edge, the half-edges that it interacts with on its tail and on its head are designated as *previous-h* and *next-h* respectively. Half-edges that are open (have no interaction partner) on their head and/or tail are designated as *boundary half-edges*, and drawn as red arrows. Each boundary half-edge has a *next-boundary-h* and a prev-boundary-h, but there is not necessarily an interaction between a half-edge and its *next-boundary-h* or *prev-boundary-h*. **(B)** The *vertex move*, in which a vertex is randomly displaced. **(C)** *Single edge addition/removal*. The single edge addition move adds an edge (two half-edges) to the boundary of the shell, if the two selected boundary half-edges are bound to each other. This results in two new interactions, drawn as dark gray wedges in the schematic. The reverse move, a single edge removal, breaks two interactions and removes two half-edges form the shell. **(D)** The *Paired edge addition/removal* move adds/removes two edges to the shell, resulting in forming/breaking three interactions (dark gray wedges in the schematic). **(E)** The *Simple boundary bind/unbind* makes/breaks an interaction between two boundary half-edges. **(F)** *Wedge fusion bind / wedge fission unbind*. A wedge fusion move adds an interaction between two boundary half edges that are sufficiently near each other. In the example shown, in the left configuration there are two nearby boundary half-edges that can bind. This causes a vertex to be removed and adds two new interactions. The reverse move, wedge fission unbinding, results in adding a new vertex and breaking two interactions. **(G)** *Fusion bind /fission unbind*. In fusion binding, two close boundary half-edges are bound, resulting in removal of a vertex and one new interaction (dark grey wedge in the schematic). The reverse move, fission unbinding, results in removal of one vertex and breakage of one interaction. **(H)** The *Conformational switch* move changes the conformations of the two half-edges within one edge.

### MC moves

The set of MC moves and their acceptance criteria are described here.

#### Notation

Figure S2 (A) shows a schematic of an example structure, with explanations of some notation that will be used in the following descriptions of the MC moves. Edges which have their full complement of four interactions are denoted as *non-boundary edges*. The two half-edges within a *non-boundary edge* are denoted as *non-boundary half-edges* (black arrows in Figure S2 (A)). Each non-boundary half-edge interacts with a half-edge on each of the two neighboring edges, which are denoted as *next-h* (from its head) and *prev-h* (from its tail). An example of a half-edge **h** with its next-h and prev-h colored in blue is shown in Figure S2 (A).

**FIG. S3.**
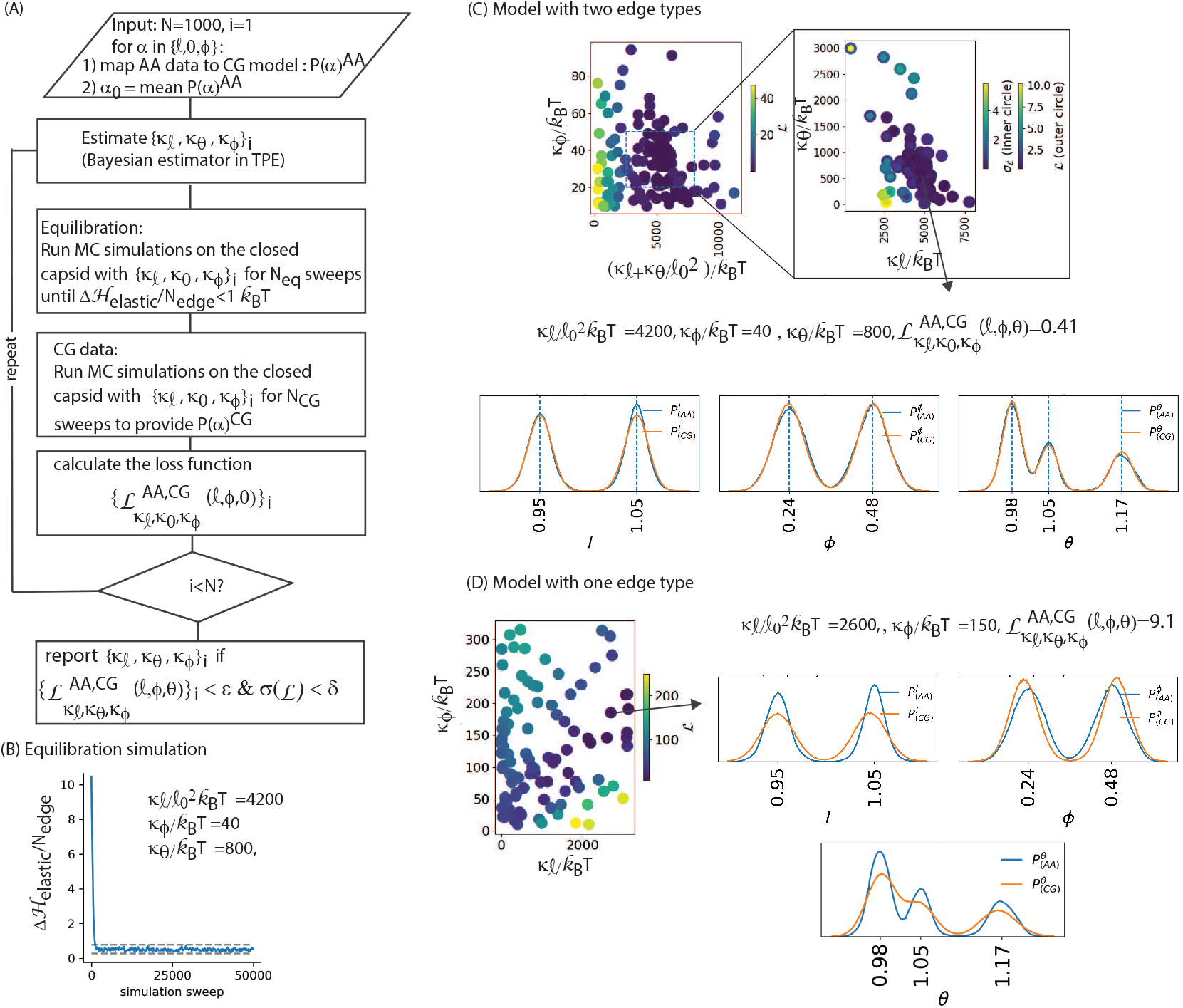
**(A)** Flow chart of parameter optimization search with the TPE algorithm of Hyperopt [152]. **(B)** Example of energy fluctuations in the equlilibration phase of simulations. **(C)** Parameter optimization results for our model with two edge types. The 3D search of *κ*_l_, *κ_θ_*, and *κ_ϕ_* is mapped to a 2D plot on the left. The zoomed-in diagram on the right shows the search as a function of *κ*_l_ and *κ_θ_*, for 20 < *κ_ϕ_* < 50. The distributions of the three parameters 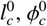, and 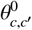 at optimized values of bending moduli are shown against the AA distribution on the bottom. Optimal values for the stretching, dihedral angle, and binding angle moduli are *κ*_l_/*σ*^**2**^*k*_B_*T* = 4200, *κ_θ_/kt* = 40, and *κ_ϕ_*/*k*_B_*T* = 800, where 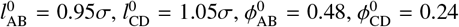, and all 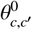 are as listed in table S1. **(B)** Parameter optimization search with the TPE algorithm for a model with only one edge type in the capsid. The optimal values and distribution of parameters are shown at right. The minimized value of the loss function is an order of magnitude larger than for the two-edge type model.

A *boundary half-edge* has at least one unbound end; i.e., it lacks an interaction with at least one of its adjoining half-edges (next-h or prev-h). By keeping track of the set of boundary half-edges, the simulation algorithm is able to efficiently choose possible trial moves which involve binding new edges. In the MC implementation used for this work, we allow only one half-edge in each edge to be a boundary half-edge, which prevents formation of dangling edges (that have only one interaction) and star-like configurations. Any trial move that results in formation of an edge that comprises two boundary half-edges is rejected.

In the description of the MC moves that follows we describe moves in terms of changes in half-edges. However, note that in general each move affects both half-edges within an edge, and these effects occur simultaneously.

##### Vertex move

A vertex is randomly chosen, and a trial is made to displace it by the vector 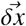, whose components are chosen from a Gaussian distribution 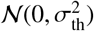. *σ*th is the length scale for the thermal fluctuations of the system which is estimated from the maximum of the three thermal lengths 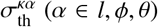 associated with bending and stretching moduli and defined above. In this move, the numbers of dimers, dimer-dimer interactions, vertices and edge types are unchanged (Fig S2(B)).

The move is accepted with probability 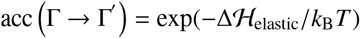.

##### Single edge addition/removal

A half-edge **h**_i_ is randomly chosen from the boundary half-edges. If **h**_i_ is bound to another half-edge **h**_j_ from one end, a trial is made to add a new edge, consisting of two half-edges **h**_k_ (and its opposite half-edge) connecting the other end of **h**_i_ and **h**_j_ (Fig S2 (C)). The generation probability is

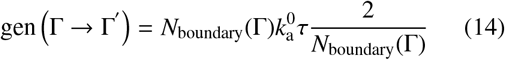

where *N*_boundary_ is the number of boundary half-edges, 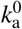 is the trial-move rate for addition/removal moves, and *τ* is the simulation time-scale explained below.

The newly added half-edge **h**_k_ interacts with **h**_i_ and **h**_j_, with conformation-dependent binding affinities, so the grand canonical probability of the new configuration relative to Γ is

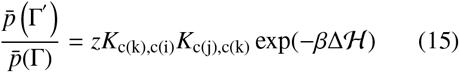

where *c*(*i*), *c*(*j*), and *c*(*k*) are the conformations of halfedges **h**_i_, **h**_j_, and **h**_k_ respectively and 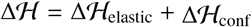 involves volume exclusion, the elastic energy of the new edge and its bound edges, and the conformational free energy of the new edge.

For removal of a single edge, a half-edge **h**_r_ is randomly chosen from the boundary half-edges. A trial is made to remove **h**_r_ and its opposite half-edge **h**_k_, which includes unbinding **h**_k_ from its next-h **h**_i_ and prev-h **h**_j_. The generation probability of edge removal is:

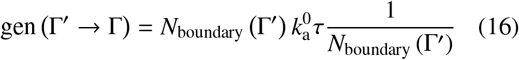

The acceptance criteria satisfying the detailed balance condition (Eq. of the main text) is:

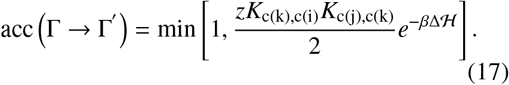

##### Paired edge addition/removal

In our model, edge additions that result in configurations with a dangling edge (which has only one interaction) are followed by addition of a second edge that closes the triangle. This choice is made because, under productive assembly conditions, dangling edges are highly unstable and quickly dissociate. Thus, simulations would spend the majority of their time on consecutive additions and removals of dangling edges. In previous Brownian dynamics simulations [65] we observed that in such situations net growth of assemblies was usually associated with either association of oligomers, or the rapid succession of additions of more than one subunit [65]. A similar conclusion was made from kinetic MC simulations [158]. To allow for this possibility, we include a move that enables additions and removals of dimers-of-dimers.

The two consecutive edge additions are attempted as follows: A half-edge **h**_i_ is randomly chosen from the boundary half-edges. If it is open on both ends (i.e. it has neither next-h nor prev-h), a trial is made to add two new edges that bind to the selected edge. This move also includes adding a new vertex to the shell, at the intersection of the two newly added edges. To specify the coordinate of the new vertex, first the equilibrium position of the new vertex 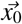 is selected based on the conformations of **h**_i_, **h**_j_ and **h**_k_. The new vertex is then displaced to 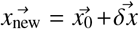 where the components of 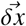 are selected from the Gaussian distribution 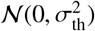 with *σ*th described above. The generation probability is

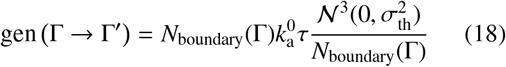

where 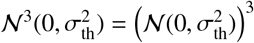.

The first added half-edge (**h**_j_) makes a new interaction with one end of **h**_i_, and the second added half-edge (**h**_k_) interacts with the open ends of **h**_j_ and **h**_i_.

For the reverse move, a a half-edge **h**_r_ is randomly chosen from the boundary half-edges. If removal of this half-edge (and its opposite half-edge) results in a dangling edge, the dangling edge will also be removed. In this move, two edges, three interactions, and a vertex are removed. The generation probability for removal

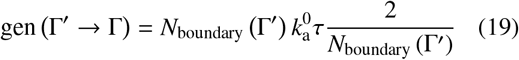

and the acceptance criteria for addition is:

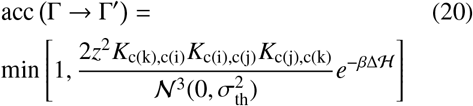

where again 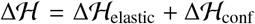 involves volume exclusion, the elastic energy of the new edges and their bound edges, and the conformational free energy of the new edges.

##### Simple boundary binding/unbinding

In a simple boundary binding, we attempt to make a new interaction between two edges whose ends are nearby but unbound to each other (Fig. S2E). For this move, a halfedge **h**_i_ is randomly chosen from the boundary half-edges. If **h**_j_ is open on both ends (has neither next-h nor prev-h) and makes a wedge with the next-boundary-h or prev-boundary-h **h**_j_, (i.e., the angle between the two edges *θ* < *π*/2) an attempt is made to bind **h**_j_ to **h**_j_.

For the opposite process, simple boundary unbinding, an edge is chosen randomly from the boundary edges and if it has a next-h or prev-h (it cannot have both since it is a boundary edge), an attempt is made to unbind. The acceptance criteria for simple boundary binding is:

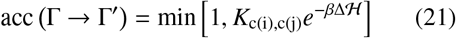

Here 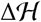 only involves the excluded volume and elastic energies.

##### Wedge fusion binding/Wedge fission unbinding

For a wedge fusion move, a half-edge **h**_i_ is randomly chosen from the boundary half-edges. A wedge fusion is attempted if: 1) **h**_i_ makes a wedge with angle *α* < *π*/2 with another boundary half-edge **h**_j_, 2) *v*_in_ of **h**_j_ is within *δx*_fuse_ of *v*_out_ of **h**_j_, 3) the next-boundary-h of **h**_i_ (**h**_k_ in Fig S2(F)) is the same as prev-boundary-h of **h**_j_, and 4) **h**_k_ is bound to **h**_j_ or **h**_i_. For the implementation of this attempt, **v**_i_ and **v**_j_ and their associated edges are fused to a new vertex **v**_k_ at the midpoint between **v**_i_ and **v**_j_. Two new interactions are made: **h**_i_ is bound to **h**_j_ and the third half-edge in the triangle **h**_k_ is bound to **h**_i_ (or to **h**_j_).

For a wedge fission unbinding, a vertex **v**_k_ is chosen at one end of a randomly chosen boundary half-edge. An interaction of that vertex between an associated half-edge **h**_i_ and its next-h **h**_j_ is randomly chosen for the wedge fission unbinding attempt. Then, *v*_out_ of the incoming half-edge **h**_i_, with all its associated edges, is moved to the new vertex **v**_i_ at 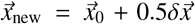; and *v*_in_ of the outgoing half-edge **h**_j_, with all its associated edges, is moved to the new vertex **v**_j_ at 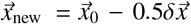, where the components of 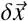 are chosen from the Gaussian distribution 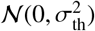. To maintain the proper topology of the shell, the interaction of the third half-edge in the triangle (randomly chosen as **h**_k_ - **h**_i_ or **h**_k_-**h**_j_) is also removed. The acceptance criteria for wedge fusion binding is:

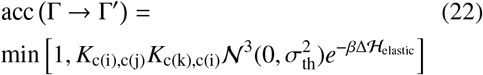

##### Fusion binding/fission unbinding

For a fusion binding move, a half-edge **h**_i_ is randomly chosen from the boundary half-edges. The move is attempted if: 1) **h**_j_ forms a wedge with angle *α* < *π*/2 with another boundary half-edge **h**_j_, 2) *v*_in_ of **h**_j_ is within *δx*_fuse_ of vout of **h**_i_, and 3) the next-boundary-h of **h**_i_ (**h**_k_ in Fig S2(G)) is not the same as prev-boundary of **h**_j_. Similar to the wedge binding move, two vertices and their associated edges are fused into a new vertex **v**_k_ at the midpoint between **v**_i_ and **v**_j_. This moves results in an additional boundary loop in the structure (orange loop in Fig S2(G) right) and the bound vertex will be a double-boundary vertex, meaning that it is shared between two boundary loops. A boundary loop can be found by starting from a random boundary half-edge, and moving along the next-boundary-h elements until returning to the original half-edge.

A fission unbinding move is attempted if there is more than one boundary loop in the structure. A vertex **v**_k_ is chosen at one end of a randomly chosen boundary half-edge. If this vertex is a double-boundary vertex, the fission unbinding move will be attempted, by splitting the edges ending in **v**_k_, to form to vertices **v**_i_ and **v**_j_ (similar to wedge fission unbinding move), and merging the two boundary loops. The implementation of adding the new vertex to the shell is similar to the wedge fission unbinding move, except that there is only one unbinding event (**h**_i_-**h**_j_).

The acceptance criteria for fusion binding is:

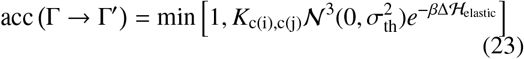

##### Conformational switch

A half-edge **h**_j_ is randomly chosen from the set of all edges in the structure, and trial is made to change the conformation of **h**_j_ from *c*(*i*) to *c*′(*i*) and the conformation of its opposite halfedge **h**_j_ from *c*(*j*) to *c*′(*j*). The move is accepted with probability:

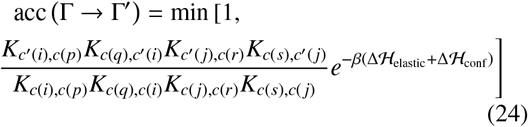

where *p* and *q* are the indices of the next-h and prev-h of *i*, and *r* and *s* are the indices of the next-h and prev-h of *j*.

### Simulation timescale

We can estimate the simulation timescale *τ*, in units of 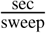, by comparing the elongation rate constants in simulations *f*_sw_ and the duration of the lag-phase *t*_lag_ before assembly occurs in experiments. This comparison is based on the theoretical finding that the lag-phase duration is proportional to the elongation timescale [159], which was experimentally confirmed by Selzer et al. [160]. Since our simulations start from the trimer-of-dimers (the critical nucleus), the *τ* does not include the nucleation time and can be used to estimate the elongation rate constant in simulations. Using the elongation rate constant calculated in HBV capsid assembly experiments ≈ 2 × 10^7^ M^-1^ s^-1^ [160] and the rate constant calculated from simulation trajectories of Fig. 3 main text. 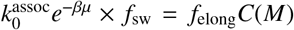 where 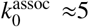, we estimate the simulation timescale *τ* ≈ 120/*f*_sw_ ≈ 10^-5^ sec/sweep.

**FIG. S4.**
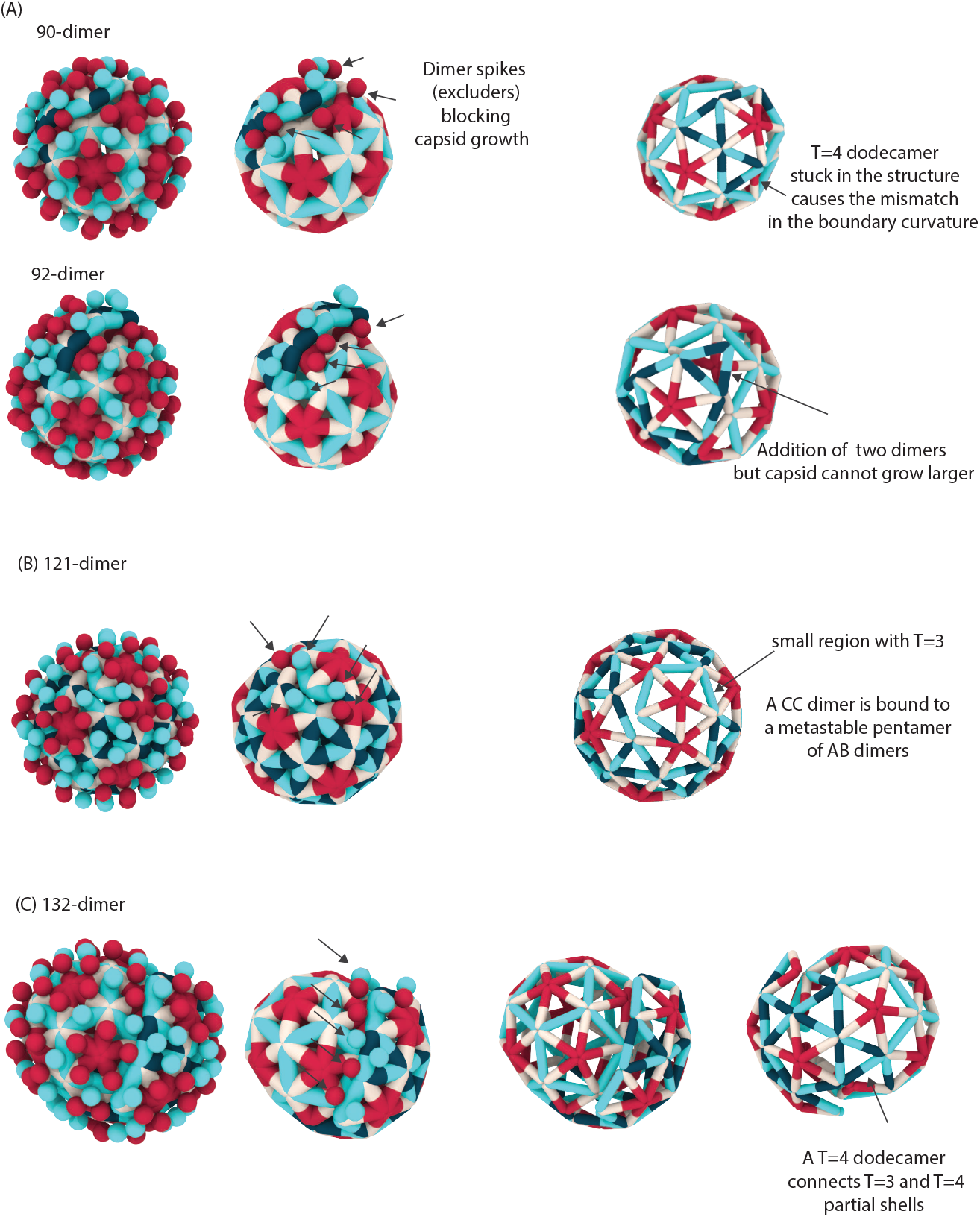
Snapshots of mixed-morphology shells. **(A)** A local *T*=4 symmetry (T=4 dodecamer) stuck in a *T*=3 intermediate results in fluctuations between 90-dimer and 92-dimer structures with open boundaries. Growth is halted due to the mismatch in the curvature, and the presence of dimer spikes (indicated by spheres and arrows in the images). **(B)** A local *T*=3 symmetry region remaining in the structure of a *T*=4 intermediate results in a long-lived holey capsid with 121 dimers. **(D)** A T4-dodecamer grows with *T*=3 symmetry from one side and *T*=4 symmetry from another side, resulting in a 132-dimer shell with mixed morphology.

**FIG. S5.**
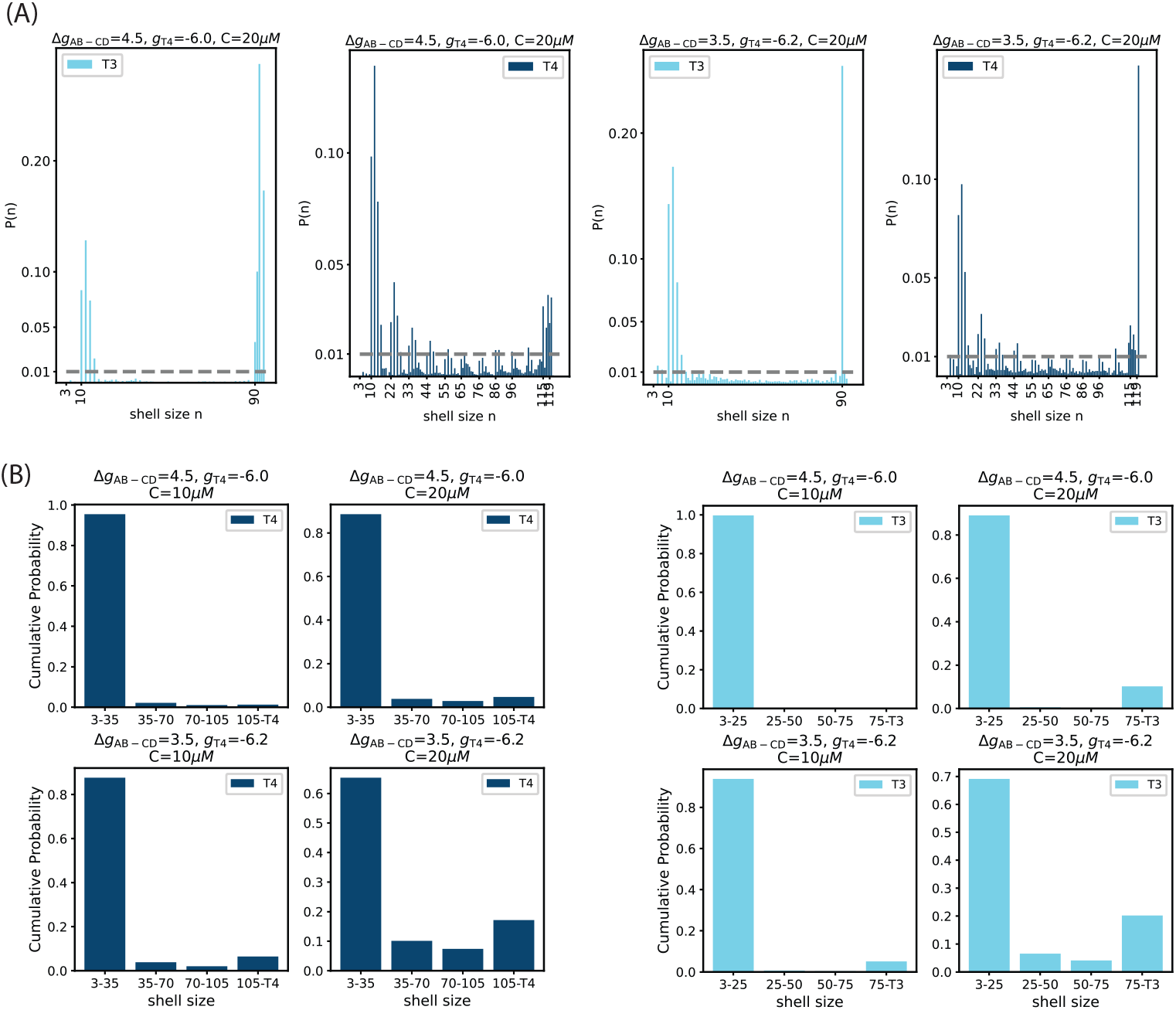
**(A)** Relative life-time of states with *n*_dimer_ > 3 in *T*=4 and *T*=3 pathways at two different parameter sets. **(B)** Cumulative time spent in different phases during the assembly of *T*=4 and T=3 capsids at selected parameter sets from the data shown in Fig. 3(A) of the main text.

## Notes

### Competing Interest Statement

The authors have declared no competing interest.

